# Lateral gene transfer shapes the distribution of nitrogen fixation within a cosmopolitan clade of marine *Thalassolituus*

**DOI:** 10.64898/2026.08.28.747955

**Authors:** Soma Sardar Barawi, Julie LaRoche, Robert G. Beiko

**Author notes:** Corresponding Author: Robert G. Beiko, Faculty of Computer Science 6050 University Avenue, Halifax, Nova Scotia, Canada B3H 4R2.

## Abstract

Biological nitrogen fixation converts dinitrogen gas into ammonia, supplying new bioavailable nitrogen to marine ecosystems, but the evolutionary processes shaping its distribution among heterotrophic bacteria remain unresolved. *Thalassolituus*, a genus within the family Oceanospirillaceae (order Oceanospirillales), is best known for hydrocarbon degradation, yet nitrogen fixation has been confirmed in only one cultured isolate. We analyzed 74 quality-filtered genomes assigned to *Thalassolituus* within a broader dataset of 421 Oceanospirillaceae genomes to reconstruct the distribution and evolutionary history of the minimal *nifHDKENB* gene set. Twenty-five genomes encoded complete or near-complete *nif* loci and occurred in four well-supported clades interspersed with genomes lacking the pathway. Statistical topology tests rejected the species-tree topology for concatenated NifHDK and NifHDKENB protein alignments, and eleven recombination events across *nif* loci were supported by at least four detection methods. The core *nifHDK* gene order remained broadly conserved, but accessory neighborhoods differed among clades, and structural *nifHDK* genes showed stronger codon adaptation than biosynthesis *nifENB* genes. Clade 2 combined species-gene tree congruence, conserved gene neighborhoods, and comparatively high *nifH* codon adaptation, whereas Clades 1 and 4 showed greater phylogenetic discordance, more recombination, and weaker codon adaptation. These results support a reticulate history in *Thalassolituus*, in which lateral acquisition introduced nitrogen fixation into distinct lineages, vertical inheritance preserved it within some clades, and homologous recombination continued to reshape *nif* loci. These processes help explain why nitrogen fixation is unevenly distributed among closely related marine heterotrophic bacteria.

## Introduction

Nitrogen (N) is a fundamental element essential for life, forming the backbone of nucleic acids, proteins, chlorophyll, and other important biomolecules (Zhang et al. 2020). Although nitrogen is abundant as dinitrogen (N₂) gas, this molecule’s chemically inert triple bond makes it unusable by most organisms, often making nitrogen the limiting nutrient for primary productivity across terrestrial and aquatic ecosystems. Biological N₂ fixation (BNF) partially solves this problem by converting inert N₂ gas into ammonia (NH₃), providing the only biological pathway supplying new bioavailable N into surface waters, which is critical in oligotrophic regions where nitrogen is often limited (Zehr and Capone 2020; Zehr and Capone 2021). This input is balanced by losses of bioavailable N through denitrification and anaerobic ammonium oxidation (anammox) processes, which return nitrogen back to the atmosphere as N₂ gas (Zhang et al. 2020). Nitrogen fixation is an energetically demanding pathway carried out by diverse prokaryotes, collectively known as diazotrophs, that possess the enzyme nitrogenase (Zhang et al. 2020; Zehr and Capone 2021). The molybdenum-dependent (Mo) nitrogenase is the most common form, encoded by the core genes *nifH*, *nifD*, and *nifK*, which together encode the dinitrogenase reductase (Fe protein) and dinitrogenase (MoFe protein) subunits (Seefeldt et al. 2020). Additional accessory genes, including *nifE*, *nifN*, and *nifB*, are involved in FeMo-cofactor assembly and cofactor biosynthesis. Together, these six genes are considered the minimal gene set required for nitrogen fixation (Dos Santos et al. 2012).

Historically, marine N₂ fixation was thought to be dominated by cyanobacterial diazotrophs in tropical and subtropical surface waters and was therefore primarily studied through targeted field observations and microscopy, with only a few diazotrophs available in culture. These works established the importance of groups such as *Trichodesmium*, *Crocosphaera*, and diatom-diazotroph associations in these regions (Capone et al. 2005). Molecular approaches targeting the *nifH* gene, a standard marker for diazotrophic diversity (Morando et al. 2025), revealed additional symbioses between haptophytes and cyanobacteria, including the nitrogen-fixing organelle UCYN-A (*Candidatus* Atelocyanobacterium thalassa) (Coale et al. 2024), as well as associations between diatoms and heterotrophic bacteria (Tschitschko et al. 2024). Furthermore, *nifH* metabarcoding studies revealed that much of diazotroph biodiversity resides in non-cyanobacterial diazotrophs (NCDs) (Gradoville et al. 2017). Genome-resolved metagenomic and metatranscriptomic studies have since identified a wide range of NCDs, including heterotrophic bacterial diazotrophs (HBDs), in environments once considered unfavorable for N_2_ fixation, including oxygen-minimum zones, deep waters, and polar regions (Delmont et al. 2018; Delmont et al. 2022; Turk-Kubo et al. 2023; Morando et al. 2025). Very few cultured representatives of HBDs exist, and their complex growth requirements and lifestyles are still poorly understood (Martínez-Pérez et al. 2018; Turk-Kubo et al. 2023; Rose et al. 2024). Recent estimates suggest that particle-associated HBDs may contribute approximately 10% of global marine N₂ fixation (Chakraborty et al. 2025), which further demonstrates that marine diazotrophy is more widespread and evolutionarily complex than previously recognized.

Major uncertainties remain regarding the evolutionary origins and spread of BNF within marine NCDs. Nitrogenase is thought to have originated approximately 3.2 billion years ago in anoxic environments prior to the Great Oxidation Event (Rucker and Kaçar 2024). Despite deep conservation of the core *nifH, nifD,* and *nifK* genes, which are typically co-localized within an operon, the ability to fix N_2_ is unevenly distributed across the microbial tree of life (Raymond et al. 2004; Pi et al. 2022). Large comparative-genomic surveys show that only a small, but ecologically diverse fraction of prokaryotic genomes encode a complete Nif core and that the scattered distribution of diazotrophy reflects a complex history of vertical inheritance, gene loss, and lateral gene transfer (LGT) (Raymond et al. 2004; Zehr and Capone 2021; Nichio et al. 2025). LGT occurs frequently among prokaryotes and is a recognized mechanism of adaptation and metabolic innovation (Ochman et al. 2000; Arnold et al. 2022), and in marine microbial communities, plays a major role in shaping diversity and ecological function (Brito 2021; Arnold et al. 2022). The *nif* genes themselves have undergone ancient duplications, losses, and frequent LGT, resulting in histories that often conflict with species phylogenies that carry them (Raymond et al. 2004; Zehr and Capone 2021; Pi et al. 2022; Nichio et al. 2025). Phylogenetic incongruence between *nif* gene trees and host species trees complicates efforts to understand how N₂ fixation is acquired and maintained within specific taxa. Given its high energetic cost, the sporadic yet widespread occurrence of diazotrophs and *nif* genes across diverse environments suggests that this pathway can provide a strong selective advantage driving their movement across taxa (Pi et al. 2025).

Within this context, *Thalassolituus* represents a useful system for examining how diazotrophy evolves in marine heterotrophic bacteria. The recent isolation of *Thalassolituus haligoni* sp. nov., BB40 (Rose et al. 2024) established the first experimentally confirmed diazotroph in a genus best known for hydrocarbon degradation (Yakimov et al. 2004; Dong et al. 2022), yet their capacity for N₂ fixation remains poorly resolved. Most members of the Oceanospirillaceae (order Oceanospirillales) are halophilic heterotrophs that are typically low in abundance but can become enriched in specialized niches such as algal blooms and oil-contaminated marine environments (Satomi and Fujii 2014). Given its global distribution and nitrogen-fixation rates comparable to other free-living NCDs, *T. haligoni* provides a useful system to investigate how this trait was acquired and maintained within the lineage (Rose et al. 2026). Here, we present a comparative genomic and phylogenomic analysis of 74 focal genomes assigned to *Thalassolituus* within the broader Oceanospirillaceae to examine the evolutionary history of nitrogen fixation. We ask (i) which lineages encode complete *nifHDKENB* loci, (ii) whether Nif and the *Thalassolituus* species tree are congruent, and (iii) whether *nifHDKENB* genes follow strict vertical inheritance or show signatures of LGT and site-specific homologous recombination. Using complementary parametric and non-parametric approaches (Brito 2021), we identify a reticulate evolutionary history in which some *nif* loci were acquired laterally, others were vertically retained within specific clades, and still others were subsequently remodeled by homologous recombination. Our analyses therefore support a central role for LGT in the spread and diversification of *nif* genes among these bacteria and suggest a possible association between diazotrophy and adaptation to nitrogen-limited, hydrocarbon-rich marine environments.

## Materials and methods

### Genome data acquisition and quality assessment

Genomes classified within the family Oceanospirillaceae, recently reclassified as Oceanobacteraceae by Flores-Félix et al. (2025), including the genus *Thalassolituus*, were systematically downloaded from the NCBI RefSeq and GenBank databases (accessed April 2025) using the NCBI Datasets command-line tool (v17.2.0) (O’Leary et al. 2024). To remove redundancy, overlapping accessions between RefSeq and GenBank were identified and removed. Genome quality was subsequently assessed using established standards: completeness >85%, contamination <5%, total genome size ≥3 Mb, and scaffold N50 values ≥20 kb (Bowers et al. 2017; Riesco and Trujillo 2024). Applying these filters resulted in a curated dataset of 421 Oceanospirillaceae genomes (Supplementary Table 1), which provided the broader phylogenomic context for this study. From this set, 74 focal genomes were retained comprising 16 isolate genomes from cultured strains and 58 metagenome-assembled genomes (MAGs) reconstructed from metagenomic sequencing data. The MAGs included six heterotrophic bacterial diazotroph (HBD) genomes from the Tara Oceans project (Delmont et al. 2018) and one Arctic MAG assigned to *Thalassolituus* (Rose et al. 2024; Shiozaki et al. 2023). The 74- genome dataset also included one genome classified as *Parathalassolituus penaei* G-43, which was retained for comparative purposes because it shares 94.45–94.76 % 16S rRNA gene similarity with *Thalassolituus* species (Chen et al. 2023). Assembly quality was evaluated using QUAST (v5.2.0) (Gurevich et al. 2013), and genome completeness and contamination were estimated with CheckM2 (v1.0.2) (Chklovski et al. 2023) using both the ’general’ gradient boost model and lineage-specific neural network model under default settings (Supplementary Table 2). Pairwise average nucleotide identity (ANI) was calculated using pyANI-plus (v1.0.0) with the ANIb method (Pritchard et al. 2016), and average amino acid identity (AAI) was estimated using FastAAI (v0.1.20) (Gerhardt et al. 2025) with default parameters to verify the genus-level assignments.

### Genome annotation and identification of putative diazotrophs

Predicted proteins from *Thalassolituus* were first queried against the UniProtKB/Swiss-Prot database (Bairoch and Apweiler 2000) using BLASTP (v2.14.1) (Camacho et al. 2009) with an expectation-value threshold of 1e-5, and candidate Nif protein sequences were extracted and checked against the NCBI non-redundant (nr) database with a more stringent e-value cutoff of 1e-20. All genomes were subsequently annotated with Bakta (v1.6.1) (Schwengers et al. 2021) to maximize gene recovery and functional annotation. Putative diazotrophic genomes were verified with the Diaiden pipeline (Chen et al. 2024) by screening for the minimal *nifHDKENB* gene set using Prodigal-predicted coding sequences (v2.6.3) and DIAMOND (v2.1.6) searches against a *nif*-specific KEGG database (updated 2025/10/28) with parameter settings ‘--sensitive -k 1 -e 1e- 100 --id 50 --query-cover 75 --subject-cover 75’ (Hyatt et al. 2010; Buchfink et al. 2015). Top homologous hits from BLASTP searches are summarized in Supplementary Table 3.

Hydrocarbon-degradation genes involved in alkane, aromatic, and plastic aerobic degradation were identified by querying proteins against the HADEG database (Rojas-Vargas et al. 2023) using Proteinortho v6.1.7 (Lechner et al. 2011).

### Mapping NifH sequence matches in Tara Oceans

The *nifH* nucleotide sequences from the 25 putative diazotrophs were first clustered at 100% sequence identity using CD-HIT (v4.8.1) (Li and Godzik 2006) with parameter settings ‘-c 1.0 -n 10 -d 0 -aS 0.05 -aL 0.95’. Nine representative sequences were selected from the resulting clusters, and NifH proteins were queried against the prokaryote-enriched Tara Oceans Microbiome Reference Gene Catalog Dataset with arctic data (OM_RGC_v2_meta) using BLASTP with an e-value cutoff of 1e-40, via Ocean Gene Atlas (v2.0) (Vernette et al. 2022) (https://tara-oceans.mio.osupytheas.fr/ocean-gene-atlas/). Abundance was normalized using the default ‘percent of total genes per sample’, which divides the sum of homolog abundances by the total gene abundance per sample. Geographic distribution maps across the five depth zones were generated in R (v4.4.1) using ggplot2 (v4.0.3) (Wickham 2016).

### Pangenome and phylogenomic analyses

Pangenomic and phylogenomic analyses were performed using the ARETE pipeline (https://github.com/beiko-lab/arete). Panaroo (v1.5.1) was used to infer core, accessory, and unique gene families across the *Thalassolituus* dataset (Tonkin-Hill et al. 2020). Core genes annotated by Bakta and identified by Panaroo were individually aligned with MAFFT (v7.487) (Katoh and Standley 2013) and concatenated into a supermatrix. A maximum-likelihood phylogeny was inferred using FastTree (v2.1.11) under a GTR model with CAT approximation for rate heterogeneity (Price et al. 2010). The *Thalassolituus* tree was rooted using four Oceanospirillaceae outgroup genomes: *Litoribacillus peritrichatus*, *Marinomonas mediterranea* MMB-1, *Neptunomonas concharum*, and *Marinomonas posidonica* IVIA-PO-181. A second phylogeny was reconstructed from “soft core” genes (identified via Panaroo as present in 95% to 99% of strains) shared across the *Oceanospirillaceae* dataset (n=421 genomes) using the same workflow to place *Thalassolituus* in a broader family-level context. All trees were visualized and annotated using tvBOT (v2.6.1) (Xie et al. 2023).

### Phylogenetic analysis of Nif proteins

Amino-acid sequences encoded by the minimal *nifHDKENB* gene set were extracted from genomes containing putative *nif* loci. Each protein was aligned separately with MAFFT (v7.511) with the --auto option. Protein trees were reconstructed with IQ-TREE2 (v2.3.6) (Minh et al. 2020), with the best-fitting substitution models selected using ModelFinder Plus (Kalyaanamoorthy et al. 2017) and branch support estimated with 1000 ultrafast bootstrap replicates (Hoang et al. 2018). Concatenated alignments of NifHDK and NifHDKENB were generated using AMAS (v1.0) (Borowiec 2016) to retain gene boundaries, and trees were inferred from these concatenated alignments in IQ-TREE2 using a partition file. Protein trees were rooted using the corresponding homologous proteins from *Bradyrhizobium diazoefficiens* USDA 110.

An expanded NifH phylogeny was constructed from 4550 bacterial and archaeal sequences retrieved from the NCBI RefSeq database, filtered to length thresholds of 250-350 amino acids and excluding partial sequences. *Thalassolituus* NifH sequences were added, and all proteins were aligned using MAFFT (v7.511). A maximum-likelihood phylogeny was inferred using IQ-TREE2 (v2.3.6) using the LG+G8 substitution model and 1000 ultrafast bootstrap replicates. The resulting tree was visualized using tvBOT (v2.6.1) (Supplementary Fig. 7).

### Tests of phylogenetic incongruence and codon-usage analysis

Evidence for lateral gene transfer was assessed through a combination of phylogenetic incongruence, statistical topology tests, and compositional analyses. To evaluate whether *nif* loci were congruent with the species tree, we compared tree topologies using generalized Robinson-Foulds (RF) distance metrics implemented in the TreeDist package (Smith 2020) in the R statistical language. Classical RF distances, which treat all bipartitions equally, quantify discordance by counting conflicting splits between trees, but they can be sensitive to tree balance and can saturate quickly especially when a single taxon is repositioned. For this reason, we used clustering information distance (CID), an information-theoretic generalization of RF distance that weights splits according to their phylogenetic information content (Smith 2020). Additional comparisons were computed using Quartet distances (Smith 2019). We also tested topological discordance using the Approximately Unbiased (AU) test (Shimodaira 2002), which compared site-wise log likelihoods of three topologies: unconstrained gene trees, the species tree, and the species-tree-constrained topology. Tests were performed using the resampling estimated log-likelihoods (RELL) approach with 10000 replicates (Kishino et al. 1990), and topologies with *P*- value < 0.05 were rejected. Phylogenetic incongruence was visualized with tanglegrams generated with the R package dendextend (Galili 2015).

In parallel, codon usage patterns of *nif* genes were analyzed to assess compositional signatures associated with potential LGT. Codon usage patterns across all 74 *Thalassolituus* genomes were calculated from coding DNA sequences (CDS) using the R package Cubar v1.2.0 (Liu et al. 2026). After filtering CDS with the ‘check_cds’ function, relative synonymous codon usage (RSCU) frequencies were determined for all 64 possible codons. Codon adaptation index (CAI), which measures similarity between a gene’s codon usage and that of highly expressed genes, was calculated using genome-specific reference sets of highly expressed ribosomal protein genes (Sharp and Li 1987), which reflect optimal codon usage patterns for each genome. To assess whether *nif* genes showed differential codon optimization relative to the host genome, *nif* CAI values were converted to percentile ranks within the CAI distribution of reference ribosomal genes in the corresponding genome (Supplementary Table 5). Lower percentile values indicated lower codon optimization relative to ribosomal genes and were used as supporting evidence alongside phylogenetic analyses.

### Recombination detection and sequence similarity analysis

Potential recombination within *nif* loci was assessed using the Recombination Detection Program (RDP5) (Martin et al. 2021). Analyses were performed on nucleotide alignments of the *nifHDKENB* genes after removing identical sequences and using default parameters. Events detected by at least four statistical methods were considered high confidence and investigated further (Supplementary Table 6). Similarity and bootscanning analyses were performed and visualized with SimPlot++ (Samson et al. 2022) under the Kimura two-parameter model, a 250 bp sliding window, and a 20 bp step size. A representative *nifH* recombination event using *Thalassolituus oleivorans* SRR3933287 as the reference sequence is shown in Fig. 5.

### Genomic neighborhood analysis

To examine gene organization, ten flanking genes (five upstream and five downstream) of the *nifHDK* operon were extracted and compared with non-diazotrophic genome neighborhoods. In six genomes, *nifB* was located distantly from the *nifHDK* block and was therefore extracted and analyzed separately. These genomic neighborhoods were subsequently aligned and visualized using Clinker (v0.0.31) (Gilchrist and Chooi 2021).

## Results

### Genome quality and clustering

Following quality control, 421 Oceanospirillaceae genomes were retained for family-level analysis (Supplementary Table 1), including 74 genomes assigned to *Thalassolituus* and the closely related *Parathalassolituus penaei* (Supplementary Table 2). Genomes from *Thalassolituus* isolates were consistently high quality, with completeness above 99% and contamination below 1.07%; six were single-contig assemblies (Supplementary Fig. 1). MAGs were more variable in contiguity but still passed the stated inclusion thresholds, with completeness and contamination >85% and <5%, respectively. Genome sizes ranged from 3.08 to 4.47 Mb, and contig counts ranged from 1 to 329. Pairwise ANI and AAI analyses separated the genomes into discrete clusters consistent with species-level delineation at 95% ANI (Supplementary Fig. 3). Most comparisons showed AAI values above 65%, which is consistent with reported genus-level ranges (Konstantinidis et al. 2017). However, three genomes showed lower similarity to the remainder of the dataset; *Thalassolituus* sp. 3umA1Z08042.57 showed 49- 52% AAI to other genomes, while two uncultured MAGs shared >95% AAI with each other but only 53-54% with the remaining genomes. These values are within ranges typically observed between different genera in the same family (Konstantinidis et al. 2017).

### Biogeographic distribution of representative NifH sequences

We used the Tara Oceans Microbiome Reference Gene Catalog Dataset with arctic data (OM_RGC_v2_meta) to investigate the distribution and abundance of exact matches to representative *Thalassolituus* NifH protein sequences across global ocean depth layers. Matches were required to have 100% amino acid identity across at least 288 amino acid residues, covering nearly the full length NifH protein. The number of hits, normalized abundances, sampling layers, and associated environmental parameters are reported in Supplementary Table 4. As shown in Fig. 1, the distribution and abundance of NifH protein sequence matches varied among clades and water depths. Sequences were categorized into four clades based on their phylogenetic placement in the *Thalassolituus* reference tree (detailed in the phylogenetic section and Fig. 3).

**Figure 1.**
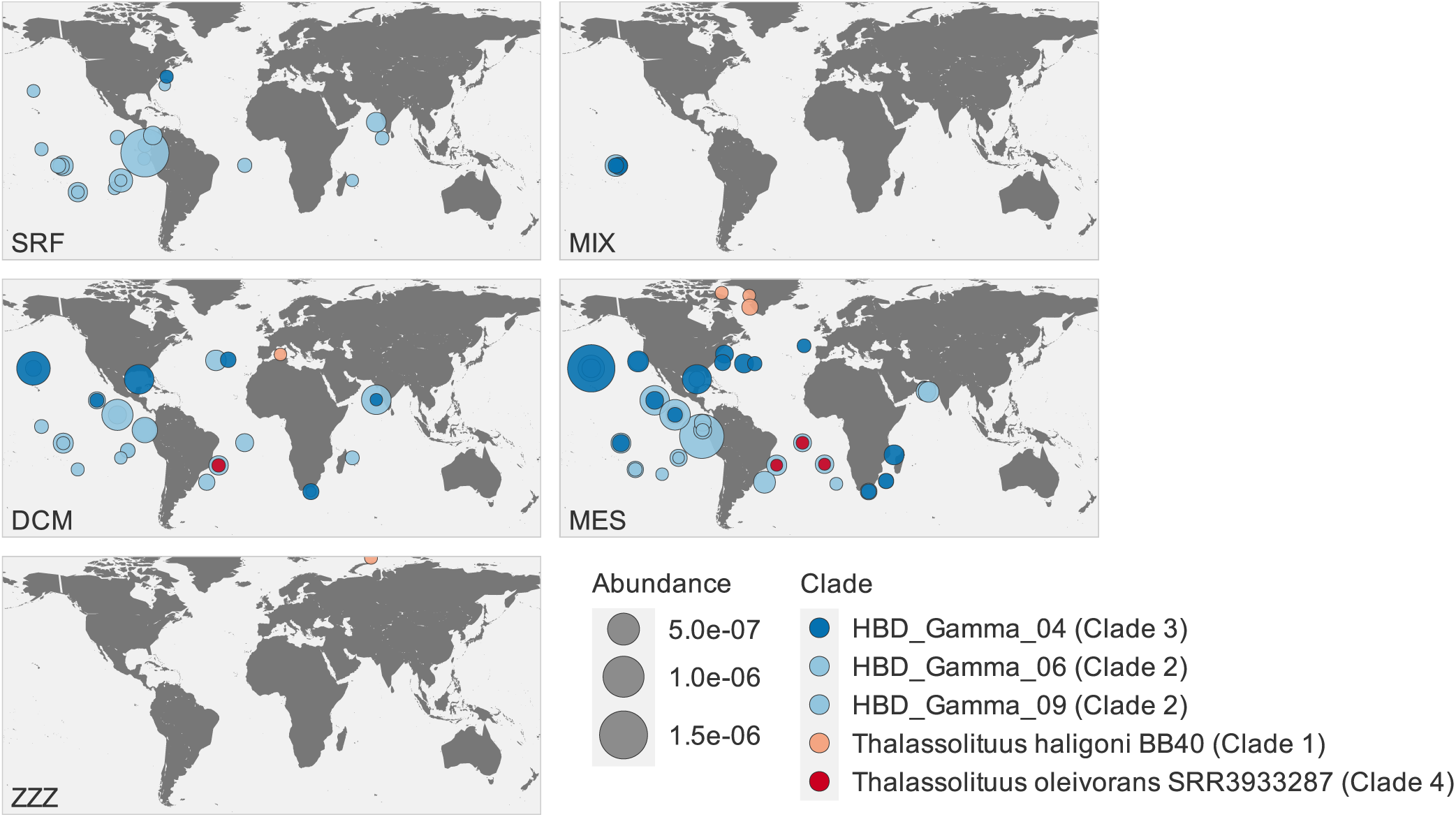
Representatives of four putative diazotrophic lineages occur across multiple ocean basins and depth zones. Normalized abundances of NifH protein matches to representatives of Clades 1-4 are shown across five depth zones: surface water layer (SRF), marine epipelagic mixed layer (MIX), deep chlorophyll maximum layer (DCM), mesopelagic zone (MES), and marine water layer (ZZZ). Matches were identified in the Tara Oceans metagenomic gene catalog (OM_RGC_v2_meta). Only matches with 100% amino acid identity across at least 288 amino acids were included.

**Figure 2.**
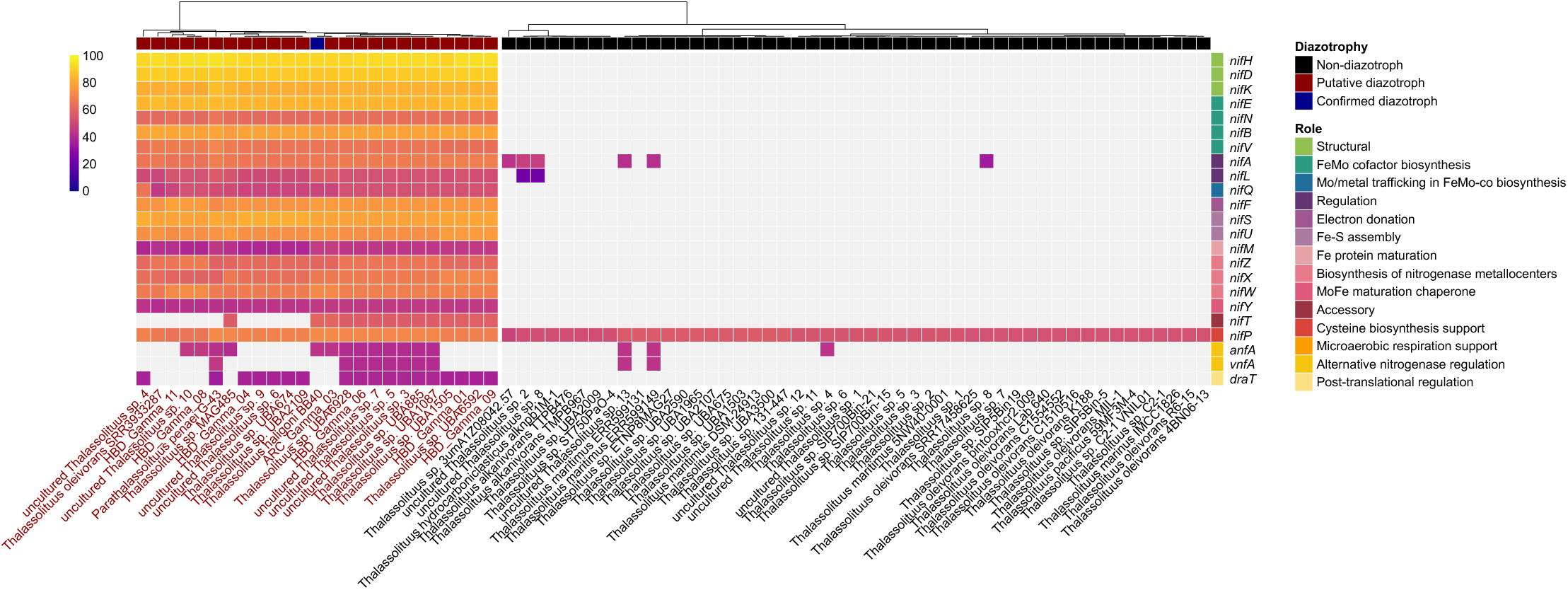
Distribution and sequence identity of Nif proteins in *Thalassolituus*. The heatmap shows the amino acid percent identity of the top BLASTP match for each Nif protein across the 74 focal genomes. Protein functional categories are indicated by the annotation colors.

**Figure 3.**
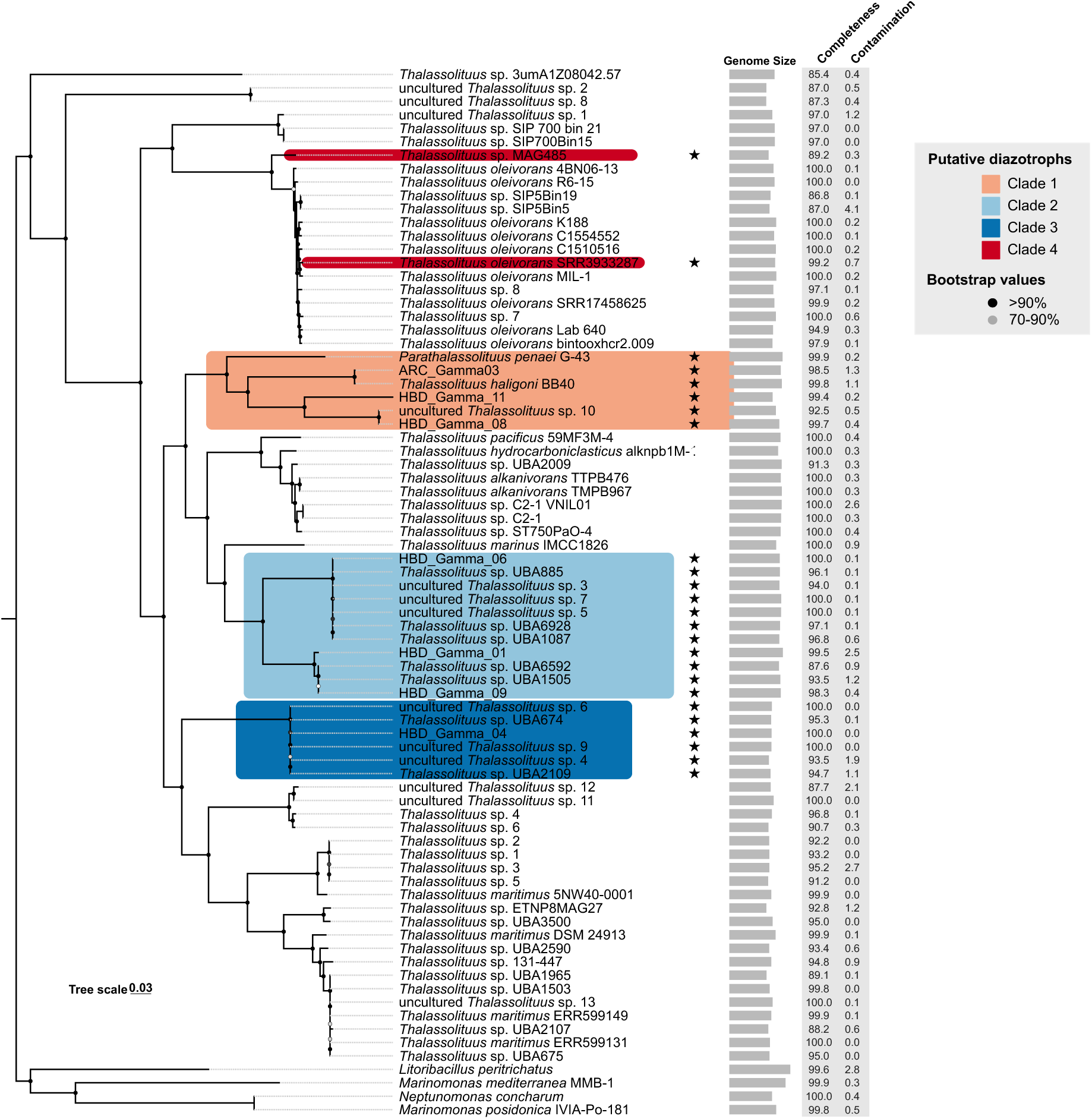
Phylogeny of *Thalassolituus*. Maximum-likelihood tree reconstructed from a concatenated alignment of 74 single-copy core genes identified across 74 *Thalassolituus* genomes and four outgroups. Putative diazotrophic genomes (n=25) are highlighted according to clade. Bootstrap support is indicated by black circles >90% and gray circles 70-90%. The scale bar represents the mean number of substitutions per site. The tree was rooted using four Oceanospirillaceae outgroups: *Litoribacillus peritrichatus*, *Marinomonas mediterranea* MMB-1, *Neptunomonas concharum*, and *Marinomonas posidonica* IVIA-PO-181.

Clade 2 sequences represented by HBD_Gamma06 and HBD_Gamma09, showed the broadest distribution with detections across multiple biogeographic provinces across the Atlantic, Indian, and Pacific Oceans and across surface (SRF), deep chlorophyll maximum (DCM), and mesopelagic (MES) layers. Clade 3, represented by HBD_Gamma04, was rarely detected in surface waters and was instead more abundant in the DCM and MES layers. Within the marine epipelagic wind mixed layer (MIX) of the South Pacific Ocean, very few NifH sequences from Clade 2 and Clade 3 were detected. Clade 1 NifH (represented by *T. haligoni)* was detected mostly at high-latitude environments in the Arctic and North Atlantic, with few matches in the DCM, MES, and ZZZ (marine water) layers. The Clade 4 representative from *T. oleivorans* SRR3933287 produced few matches, all from DCM and MES samples in the South Atlantic Ocean.

### Patchy distribution of nitrogen fixation genes

Screening predicted proteins against UniProtKB/Swiss-Prot and NR databases using BLASTP, combined with Bakta and Diaiden annotations, revealed a patchy distribution of *nif* genes across *Thalassolituus* (Fig. 2). Of 74 genomes, 25 (33.8%) encoded complete or near-complete *nif* gene sets and were classified as putative diazotrophs (Table 1), including 23 MAGs and the isolate genomes of *T. haligoni* and *P. penaei*. The remaining 49 genomes lacked the core *nifHDKENB* genes. Among putative diazotrophs, the structural nitrogenase genes *nifHDK* showed the highest conservation relative to curated Swiss-Prot sequences, with mean amino acid identities of 92.4% for *nifH*, 88.6% for *nifD*, and 82.1% for *nifK*. Genes involved in Fe-Mo cofactor biosynthesis and assembly were also consistently detected, including *nifE* (mean percent identity 83.7%), *nifB* (78.8%), and *nifN* (62.9%), whereas *nifV, nifQ, nifX,* and *nifY* showed lower sequence identity (42.3 to 65.8%). Accessory genes involved in Fe-S cluster assembly and protein maturation included *nifS, nifU, nifW, nifZ,* and *nifM*, which had mean identities ranging from 42.4 to 79.7%. The primary transcriptional regulatory genes *nifA* and *nifL* were detected in all putative diazotrophs but displayed greater sequence divergence (65.2% and 53.2%, respectively) compared to the core structural genes. The short gene *nifT* of unknown function was annotated in only 14 of 25 putative diazotrophs (averaging 58.3% identity) and annotated as a hypothetical protein in others, while the *nifP* gene involved in cysteine biosynthesis (Pratte et al. 2025) was present in all 74 genomes regardless of inferred nitrogen fixation capability (59% mean percent identity). Additionally, the *draT* gene, which regulates nitrogenase activity through the reversible ADP-ribosylation of the dinitrogenase reductase component, was detected in 18 of the 25 putative diazotrophs with a mean amino acid identity of 37.7%.

**Table 1.** Genome characteristics of the 25 putative diazotrophs included in this study.

| Accession | Genome | Completeness | Contamination | Genome Size |
| --- | --- | --- | --- | --- |
| ARC_Gamma03 | ARC_Gamma03 | 98.45 | 1.3 | 4284528 |
| GCA_002294235.1 | *Thalassolituus* sp. UBA885 | 96.14 | 0.1 | 4208961 |
| GCA_002298335.1 | *Thalassolituus* sp. UBA674 | 95.31 | 0.14 | 3488490 |
| GCA_002314145.1 | *Thalassolituus* sp. UBA1087 | 96.83 | 0.55 | 4168903 |
| GCA_002331085.1 | *Thalassolituus* sp. UBA2109 | 94.74 | 1.09 | 3439103 |
| GCA_002450785.1 | *Thalassolituus* sp. UBA6928 | 97.13 | 0.14 | 4227541 |
| GCA_913047515.1 | uncultured *Thalassolituus* sp. 3 | 94.01 | 0.06 | 4180274 |
| GCA_913052225.1 | uncultured *Thalassolituus* sp. 4 | 93.49 | 1.86 | 3294904 |
| GCA_937895325.1 | uncultured *Thalassolituus* sp. 5 | 100 | 0.11 | 4318164 |
| GCA_937902495.1 | uncultured *Thalassolituus* sp. 6 | 100 | 0 | 3549921 |
| GCA_963998775.1 | uncultured *Thalassolituus* sp. 7 | 100 | 0.11 | 4332405 |
| GCA_964007845.1 | uncultured *Thalassolituus* sp. 9 | 99.99 | 0 | 3504649 |
| GCA_964012525.1 | uncultured *Thalassolituus* sp. 10 | 92.55 | 0.49 | 3878315 |
| GCF_002324005.1 | *Thalassolituus* sp. UBA1505 | 93.54 | 1.19 | 4248465 |
| GCF_002433655.1 | *Thalassolituus* sp. UBA6592 | 87.62 | 0.93 | 4137207 |
| GCF_026626885.1 | *Parathalassolituus penaei* G-43 | 99.89 | 0.18 | 4444085 |
| GCF_034111725.1 | *Thalassolituus* sp. MAG485 | 89.24 | 0.28 | 3278734 |
| GCF_041222825.1 | *Thalassolituus haligoni* BB40 | 99.77 | 1.07 | 4376545 |
| GCF_913058015.1 | *Thalassolituus oleivorans* SRR3933287 | 99.19 | 0.65 | 3820462 |
| HBD_Gamma_01 | HBD_Gamma_01 | 99.45 | 2.52 | 4469063 |
| HBD_Gamma_04 | HBD_Gamma_04 | 99.99 | 0.01 | 3519070 |
| HBD_Gamma_06 | HBD_Gamma_06 | 99.97 | 0.08 | 4203397 |
| HBD_Gamma_08 | HBD_Gamma_08 | 99.65 | 0.38 | 4169142 |
| HBD_Gamma_09 | HBD_Gamma_09 | 98.35 | 0.43 | 4291755 |
| HBD_Gamma_11 | HBD_Gamma_11 | 99.37 | 0.24 | 3611393 |

Genomes lacking the essential *nifHDKENB* genes showed only sporadic, low-identity hits to regulatory *nif* genes. Alternative nitrogenase genes, including *anfA* (averaging 40.9% sequence identity) and *vnfA* (43.4%) for iron-only and vanadium-dependent systems, were either absent or appeared as low-identity hits in a small number of genomes. Top *nifHDKENB* homologs were mostly assigned to unclassified Pseudomonadota and Gammaproteobacteria, but best matches also included families outside of Oceanospirillaceae including Alteromonadaceae, Cellvibrionaceae, Pseudomonadaceae, Halieaceae, Ketobacteraceae, and Sedimenticolaceae (Supplementary Fig. 2; Supplementary Table 3). The most frequent matches from Oceanospirillaceae included *Thalassolituus* and *Oceanobacter* alongside other marine genera such as *Pseudomonas, Pseudomaricurvus, Marinobacterium, Ketobacter,* and *Mangrovimicrobium*.

### Phylogenetic relationships among Oceanospirillaceae and *Thalassolituus*

We inferred the phylogenetic placement of *Thalassolituus* genomes by constructing a maximum-likelihood phylogenetic tree based on a core-genome alignment of 74 single-copy genes shared across all genomes and outgroups. This produced a well-supported species tree (mean bootstrap support = 95.47%) and revealed that putative diazotrophs are not monophyletic but instead distributed across four distinct clades interspersed among non-diazotrophic lineages (Fig. 3). For simplicity, we refer to these putative diazotrophic clades as Clades 1-4, each of which recovered 100% bootstrap support. Clade 1 comprises three lineages: *P. penaei* G-43 as sister to a subclade containing ARC_Gamma03 and *T. haligoni*, and a subclade of three genomes (HBD_Gamma11, HBD_Gamma08, and an uncultured *Thalassolituus* sp. 10). Clades 2 (eleven genomes) and 3 (six genomes) occurred on deeper branches and consist largely of uncultured MAGs, (UBA885, UBA6928, and UBA1087) together with HBD genomes. Clade 4 contains predominantly non-diazotrophic *Thalassolituus oleivorans* genomes with relatively short internal branch lengths.

The only putative diazotrophs in this clade are *T. oleivorans* SRR3933287, which is nested within the *T. oleivorans* clade, and *Thalassolituus sp.* MAG485, which is sister to that clade. To place these lineages in a broader family context, a second maximum-likelihood tree including 421 Oceanospirillaceae genomes was constructed based on a core-genome alignment of 120 soft core genes, since strict core genes collapsed to zero at the family scale (Supplementary Fig. 4). This tree was also well-supported (mean bootstrap support 94.61%) and likewise showed that genomes assigned to *Thalassolituus* do not form a single family-level clade, but are interleaved with other genera including *Oceanobacter*, *Bacterioplanes*, *Bacterioplanoides*, several unclassified Oceanospirillaceae, and a genome assigned to *Venatoribacter*.

### Nif gene phylogenies

To test whether the evolutionary history of nitrogen fixation mirrors the host-genome phylogeny, we reconstructed maximum-likelihood trees for each of the six essential Nif proteins, and for concatenated alignments of NifHDK and NifHDKENB. Average bootstrap support values varied noticeably across individual trees, with NifH and NifD displaying the lowest average support at 63.2% and 66.3%, followed by NifB (74.7%), NifE (81%), NifK (86.3%) and NifN (89.3%).

Concatenated alignments improved resolution, with averages of 84.8% for NifHDK and 87.6% for NifHDKENB. The NifHDKENB gene tree recovered a few clades with ≥ 90% bootstrap support that were also present in the species tree, indicating a consistent phylogenetic backbone (Fig. 4A). Specifically, Clade 2 (comprising the seven-genome HBD_Gamma06 subclade and the four-genome HBD_Gamma09 subclade) was recovered with 100% bootstrap support and was congruent between both trees. However, despite this backbone, multiple topological conflicts were evident between the NifHDKENB and species trees, which we visualized using tanglegrams (Fig. 4B).

**Figure 4.**
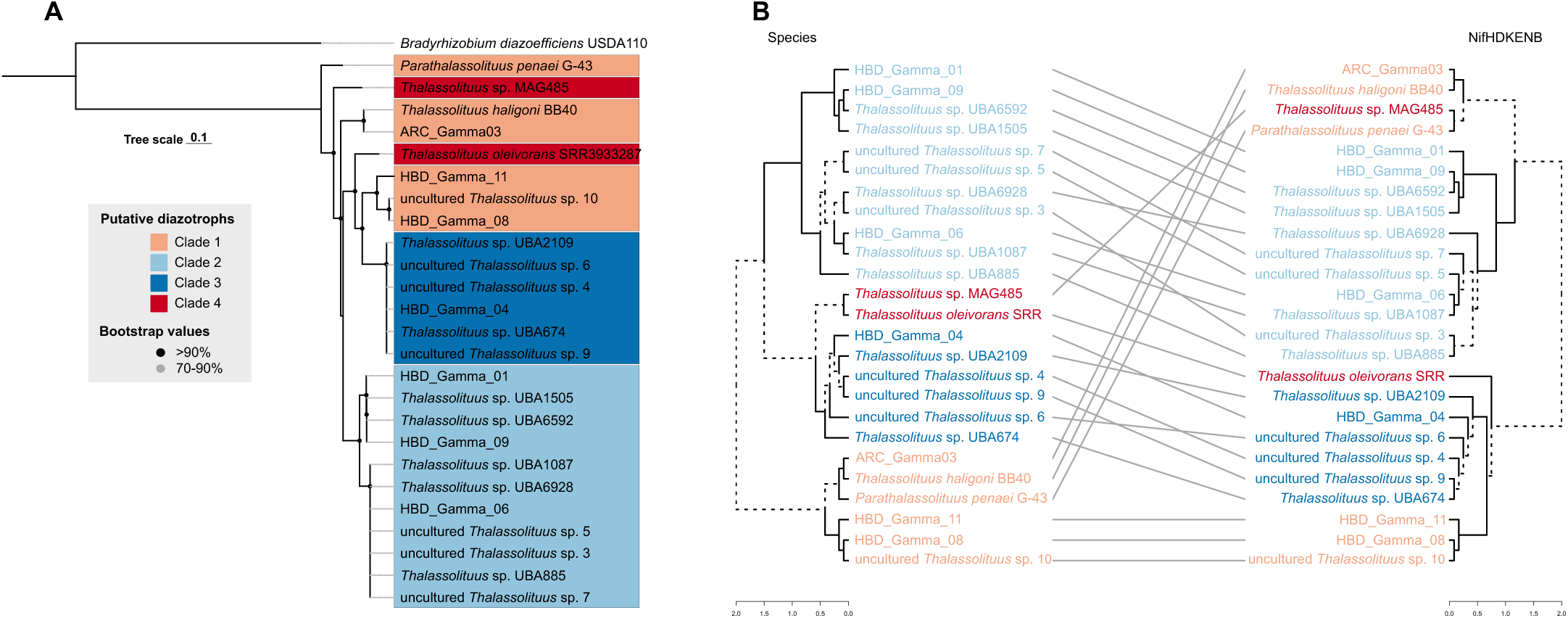
NifHDKENB phylogeny and comparison with the *Thalassolituus* reference phylogeny. (A) Maximum-likelihood tree reconstructed from a concatenated alignment of NifHDKENB protein sequences identified in the 25 putative diazotrophic genomes. Bootstrap support is indicated by black circles (>90%) and gray circles (70-90%). The scale bar represents the mean number of substitutions per site. The tree was rooted using homologous sequences from *Bradyrhizobium diazoefficiens* USDA 110. (B) Tanglegram comparing the *Thalassolituus* reference phylogeny with the concatenated NifHDKENB protein tree. Connecting lines link corresponding genomes between the trees, and dashed branches indicate edges unique to either topology.

**Figure 5.**
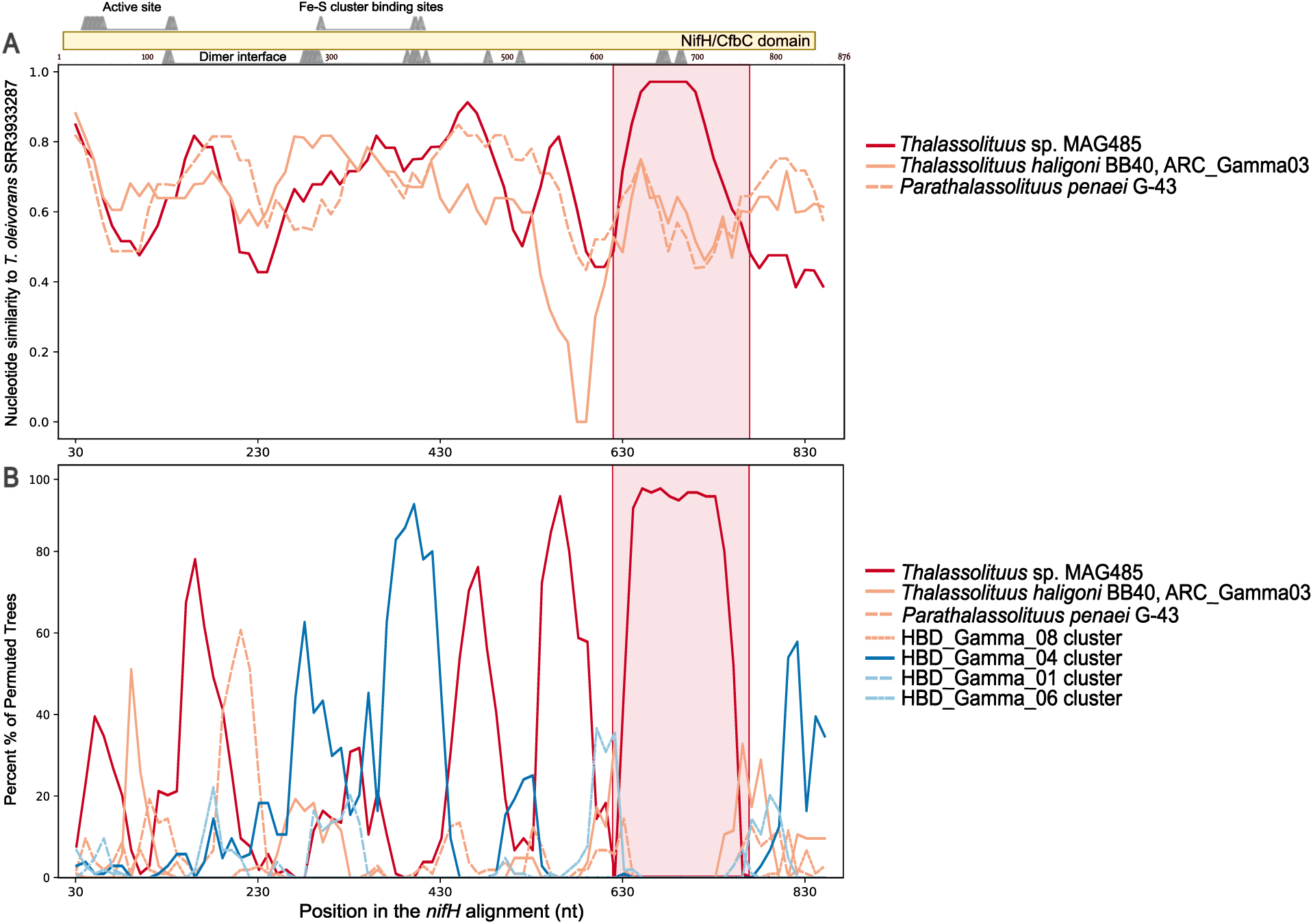
Recombination analysis of *nifH* using *Thalassolituus oleivorans* SRR3933287 as the reference sequence. (A) Sliding window analysis showing nucleotide similarity between the reference and selected sequences using a 250 bp window and 20-bp step. (B) Bootscanning analysis showing the percentage of permuted trees supporting the grouping of the reference with each comparison sequence across the alignment. The shaded region indicates the RDP5- identified recombination interval at positions 625-746. Colors represent putative diazotrophic clades, while line types distinguish sequences or clusters within the same clade.

In the species tree, Clade 1 forms a single, well-supported cluster, whereas in the concatenated NifHDKENB protein tree, its genomes are split into distinct clades. Most notably, the three-genome subclade from Clade 1 (HBD_Gamma11, HBD_Gamma08, and an uncultured *Thalassolituus* sp. 10) clusters with Clade 3 genomes to form a larger, strongly supported clade (98.4% bootstrap support) despite being separated in the species tree. Additional conflicts involve the positions of *P. penaei* G-43, *T. haligoni* and ARC_Gamma03 from Clade 1, and *Thalassolituus sp*. MAG485 and *T. oleivorans* SRR3933287 from Clade 4. In the NifHDKENB tree, *Thalassolituus sp*. MAG485 and *P. penaei* G-43 both branch early and are separated from the remaining ingroup, whereas *T. oleivorans* is recovered as sister to the large clade that includes the expanded Clade 1 and Clade 3 group, with 100% bootstrap support. We also reconstructed an expanded maximum-likelihood phylogeny of 4550 bacterial and archaeal NifH sequences from NCBI RefSeq (Supplementary Fig. 7). While this tree expectedly placed *Thalassolituus* within other Gammaproteobacteria, putative diazotrophs similarly were not monophyletic, which mirrors the topological discordance observed between the reference and Nif protein trees. Clade 3 remained largely preserved, but Clades 1, 2, and 4 were interspersed with several other genera, including *Oceanobacter, Ketobacter, Teredinibacter,* and *Marinobacterium* with varying bootstrap support values ≥70%.

Individual Nif gene trees broadly recover similar groupings of genomes as the concatenated tree, but the placement of specific taxa also varies greatly among loci (Supplementary Fig. 5). For example, the NifH tree shows more pronounced conflicts with the reference than does the NifHDKENB tree. Clade 1 genomes were split into at least three distinct lineages. *P. penaei* emerged as the earliest diverging branch of the ingroup, HBD_Gamma11 became sister to the *T. haligoni* and ARC_Gamma03 pair, and the HBD_Gamma08 and uncultured *Thalassolituus* sp. 10 pair clustered with the seven-genome subclade of Clade 2. In contrast to its congruence in the concatenated NifHDKENB and species trees, Clade 2 was fragmented in the NifH tree. The four-genome subclade of Clade 2 formed a sister group to Clade 3, with *Thalassolituus* sp. MAG485 and *T. oleivorans* also appearing within this larger group. These qualitative differences between the Nif and species tree point to systematic phylogenetic discordance that we next quantified using tree-distance and topology tests.

### Phylogenetic discordance and statistical support for species-gene tree conflicts

We quantified topological differences between the Nif protein trees and the *Thalassolituus* species tree using two complementary distance metrics. Clustering information distance (CID) is an information-theoretic generalization of the Robinson-Foulds framework that measures differences in the clustering structure of two trees, while quartet distance (QD) measures the proportion of four-taxon subsets that differ between trees. Because each resolved quartet can have one of three possible topologies, the expected QD value between two random trees is approximately two-thirds. In both cases, a value of 0 indicates identical topologies (and complete congruence with the species tree), whereas higher values indicate increasing discordance.

All Nif trees differed from the species tree, with observed CID values ranging from 9.05 to 13.06 bits (Table 2). The concatenated NifHDK and NifHDKENB trees had the lowest CID and were therefore less discordant than the individual protein trees, whereas NifH and NifN had the highest values. Similarly, QD values showed that 18.3 to 25.5% of quartets differed between the Nif and species trees. The Nif trees therefore share substantial phylogenetic signals with the species tree but are not topologically identical to it.

**Table 2.** Phylogenetic discordance and topology tests comparing Nif protein trees with the *Thalassolituus* species tree.

| Alignment | CID (bits) | QD | AU *P* value | Species-tree topology |
| --- | --- | --- | --- | --- |
| NifH | 12.53 | 0.255 | 0.212 | Not rejected |
| NifD | 11.29 | 0.21 | 0.143 | Not rejected |
| NifK | 10.82 | 0.183 | $2.64 \times 10^{-5}$ | Rejected |
| NifE | 11.77 | 0.201 | $7.77 \times 10^{-5}$ | Rejected |
| NifN | 13.06 | 0.213 | $1.16 \times 10^{-8}$ | Rejected |
| NifB | 11.98 | 0.188 | 0.00393 | Rejected |
| NifHDK | 9.85 | 0.184 | 0.00427 | Rejected |
| NifHDKENB | 9.05 | 0.195 | $1.77 \times 10^{-86}$ | Rejected |

**Abbreviations:** CID, clustering information distance; QD, quartet distance; AU, approximately unbiased test.

Approximately Unbiased (AU) tests were then used to evaluate whether each Nif alignment supported or rejected the species tree topology (Table 2). The species tree topology was rejected for most loci and concatenated alignments with *P*-values < 0.05, but not for NifH (*P*-AU = 0.212) and NifD (*P*-AU = 0.143). Failure to reject the species tree topology for these genes does not demonstrate congruence or strict vertical inheritance, rather, their high sequence conservation may limit phylogenetic signal, leaving insufficient evidence to distinguish the species tree topology from alternative topologies. Both concatenated alignments of NifHDK (*P*-AU = 0.00429) and NifHDKENB (*P*-AU = 1.77E-86) strongly rejected the species tree topology, providing the statistical support for the incongruence observed in the gene trees.

### Recombination dynamics in *nif* genes

Homologous recombination within *nif* loci is a plausible driver of the phylogenetic discordance observed in our tree topologies. To test this, nucleotide sequences were analyzed using the RDP5 program. Of the 42 putative recombination events detected across the six genes, 11 events were supported by ≥ 4 methods (Supplementary Table 6). These events repeatedly involved the same subset of putative diazotrophic genomes, primarily from Clades 1 and 4, which exhibited the most pronounced topological conflicts in our tanglegrams and acted interchangeably as putative recombinant and parental lineages. In *nifH*, three supported events were identified. The strongest occurred in *T. oleivorans* SRR3933287 (Clade 4), spanning breakpoint positions 128-624 and supported by six methods, with *Thalassolituus* sp. MAG485 (also Clade 4) inferred as the minor parent. Additional *nifH* recombination events in *T. oleivorans* SRR3933287 (positions 625 to 746) involved *T. haligoni* (Clade 1) as the major parent and *Thalassolituus* sp. MAG485 (Clade 4) as the minor parent. A sliding window similarity analysis (Fig. 5A) using *T. oleivorans* SRR3933287 *nifH* as the reference sequence showed an increase in nucleotide identity (100%) with *Thalassolituus* sp. MAG485 across this region. Bootscanning analysis revealed that within the same region, MAG485 was the nearest neighbor in approximately 100% of permuted trees, whereas outside of this window it shifted to alternative lineages, which may indicate additional recombination between sequences (Fig. 5B). The final *nifH* event, detected in *Thalassolituus* sp. MAG485 (positions 746 to 873), assigned *T. oleivorans* SRR3933287 as major parent, and *P. penaei* (Clade 1) as the minor parent.

This cross-clade exchange was also observed in other loci. In *nifD*, a single event in *Thalassolituus* sp. MAG485 (positions 1002-1210) assigned Clade 1 members HBD_Gamma11 and *P. penaei* as major and minor parents, respectively. In *nifK*, four events were identified among the same Clade 1 genomes, including *P. penaei* (positions 937 to 1062), *T. haligoni* (positions 115-656), and HBD_Gamma11 (positions 770-842). Despite support from multiple methods, exact parental assignments for these events remained uncertain. In *nifE*, a single event in *T. haligoni* (Clade 1) spanning positions 559-635 assigned HBD_Gamma08 (Clade 1) as the major parent and *Thalassolituus* sp. UBA885 (Clade 2) as the minor parent. Finally, in *nifN* and *nifB*, additional events were detected within Clade 4 genomes *T. oleivorans* SRR3933287 (positions 438-833 and 458-986), and *Thalassolituus* sp. MAG485 (positions 993-1070), though their parental lineages remained uncertain.

### Codon usage of *nif* genes

To evaluate whether *nif* genes exhibit codon usage patterns distinct from their genomic background, we examined codon usage bias in the six essential *nif* genes across the 25 putative diazotrophic genomes. To assess adaptation to the host’s translational machinery, CAI was calculated per genome using genome-specific ribosomal protein genes as the highly expressed reference set. Structural *nifHDK* genes consistently had higher mean CAI values (0.804, 0.787, and 0.790) than *nifE*, *nifN*, and *nifB* (0.647, 0.59, 0.679) (Supplementary Fig. 8; Supplementary Table 5). To allow standardized comparisons across genomes with differing baseline compositions, we converted *nif* CAI values to empirical percentile ranks relative to each genome’s specific ribosomal reference distribution. Overall, *nifHDK* genes typically occupied the upper portion of the ribosomal CAI distribution. Among these, *nifH* showed the highest relative CAI with a mean percentile rank of 75.5, followed by *nifK* (66.7) and *nifD* (66.4), respectively. In contrast, the biosynthesis genes *nifENB* displayed substantially lower ribosomal percentile ranks (2.5, 1.0, and 12.6 respectively), with *nifN* demonstrating a CAI lower than all ribosomal genes in 15 genomes.

Across individual genomes, a clear clade-level pattern was observed in codon adaptation of the structural *nif* genes. Clade 2 genomes, which were congruent between the species and *NifHDKENB* trees, showed the highest levels of codon adaptation overall. All but one genome (*Thalassolituus* sp. UBA6592, 58.8th percentile) had *nifH* ribosomal percentile ranks ranging from approximately 80 to 95.1. In contrast, genomes consistently associated with phylogenetic discordance and potential recombination showed reduced codon adaptation of *nif* genes compared to the overall distribution. This was evident in Clade 4 genomes *Thalassolituus sp. MAG485* and *T. oleivorans,* which showed reduced *nifH* percentile ranks of 52.9 and 53.2.

Similarly, *P. penaei* from Clade 1 exhibited the lowest *nifH* percentile rank in the dataset (38), while other Clade 1 genomes ranged from 67.3 (*T. haligoni)* to 80.8 (uncultured *Thalassolituus* sp. 10). In these genomes, *nifD* and *nifK* also showed reduced ribosomal-relative codon adaptation consistent with *nifH*.

### Genomic organization of *nif* gene clusters

The 25 putative diazotrophs shared a conserved *nifHDK* core but differed by clade-specific accessory regions (Fig. 6). The *nifH*, *nifD*, and *nifK* genes generally formed a syntenic block and were complete in all genomes except *Thalassolituus* sp. UBA1505, where *nifH* was truncated at the end of a contig and annotated as a hypothetical protein. *T. haligoni* was the only genome with a hypothetical protein within the *nifHDK* block. A SIR2-2 domain-containing protein adjacent to *nifH* occurred in 23 genomes and was absent from two truncated loci, while *nifT*, DUF6129 domain-containing proteins, *nifB*, a FeMo-cofactor biosynthesis protein, glutaredoxin, XH and Nitro-FeMo-Co domain-containing proteins, and hypothetical proteins were also common and conserved. The *nifE* and *nifN* genes were adjacent in all genomes but occurred on the same contig as *nifHDK* in only 16. In most cases, the *nifEN* pair was located approximately 25-27 kb from *nifHDK*, but the exceptions were *Thalassolituus* sp. MAG485 (Clade 4), where the pair was 7 kb away, and Clade 1 genomes ARC_Gamma03 and *T. haligoni,* where it was approximately 160 and 184 kb away, respectively. Similarly, *nifB* occurred on the core contig in 19 genomes and on a different contig in six genomes (Supplementary Fig. 6). Its median distance from *nifHDK* was 8.7 kb, except in ARC_Gamma03 and *T. haligoni* where it was approximately 18 kb away. Ankyrin-repeat proteins also occurred in 10 genomes across Clades 1, 2, and 4.

**Figure 6.**
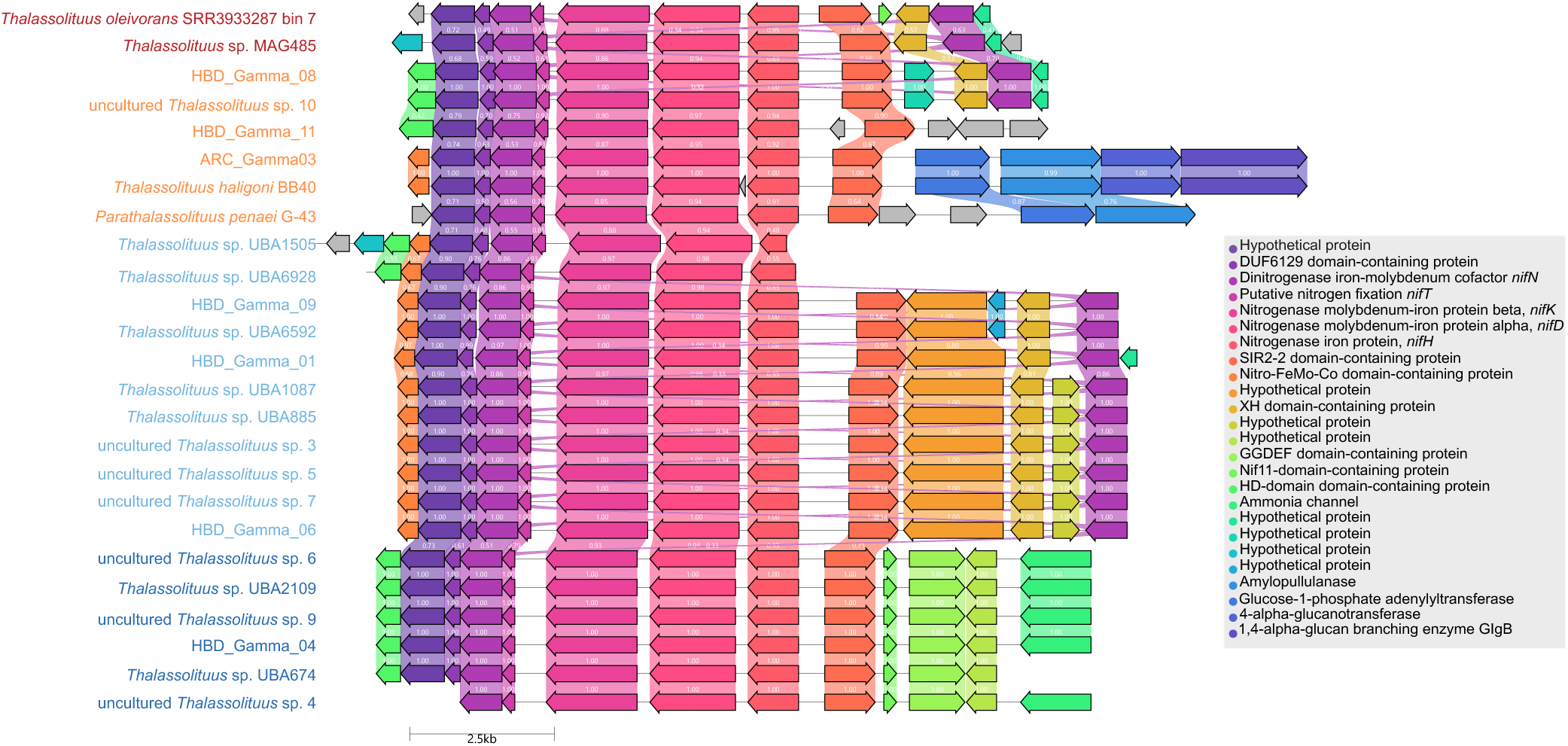
Genomic neighborhoods of *nifHDK* and ten flanking genes in 25 putative diazotrophic *Thalassolituus* genomes. Genes are drawn to scale as arrows, with their orientation indicating the direction of transcription. Colors denote homologous gene groups identified by global pairwise alignments, and connecting links represent the degree of amino acid sequence identity.

Beyond these shared genes, the genomic architecture varied largely by clade. Clade 1 showed the greatest variation in accessory gene content. *T. haligoni, P. penaei*, and ARC_Gamma03 shared genes for carbohydrate storage and energy metabolism including amylopullulanase, glycogen synthase, and glucan-associated genes, which were absent in the core *nif* neighborhoods of all the other clades. The *P. penaei* neighborhood contained unique biopolymer-transport proteins, and HBD_Gamma08 and uncultured *Thalassolituus* sp. 10 shared genes for phosphate acquisition, ion-translocating oxidoreduction, a folate-dependent homocysteine methyltransferase, a MprA stress response regulator, and an Acriflavine resistance protein. These two genomes also shared DNA topoisomerase 3 enzymes with HBD_Gamma11. HBD_Gamma08 also contained a phage-tail protein near *nifH* and a putative mobile region approximately 20-30 kb upstream of *nifHDK*, which included paired tRNA-Arg genes, a phage- associated protein, DEAD/DEAH and RecG helicases, a DNA methylase, and a site-specific integrase. HBD_Gamma11 and *P. penaei* had the most divergent *nifHDK* neighborhoods based on mean sequence similarity.

In contrast, Clade 2 showed the most conserved accessory gene organization. All 11 genomes contained a clade-specific HD-domain protein that was not detected in other clades, and nine shared a larger region downstream of *nifB* that encoded proteins involved in protein turnover, riboflavin and isoprenoid biosynthesis, response regulation, and DNA repair and recombination. These proteins were largely absent in *Thalassolituus* sp. UBA1505 and *Thalassolituus* sp. UBA6928 which contained *nifH* at contig boundaries. All six Clade 3 genomes contained a Nif11 domain-containing protein and five also carried a GGDEF domain protein.

HBD_Gamma04 uniquely had a diguanylate cyclase and a phage-tail protein near *nifHDK*. Additional genes unique to this clade near *nifHDK* included proteins with predicted roles in flavohemoglobin expression modulation, anion-transport regulation or lipid binding, ammonium transport, and cellular homeostasis. The two Clade 4 genomes shared glutaredoxin, ferredoxin- associated proteins and an ion-translocating oxidoreductase complex but differed in other accessory genes. MAG485 encoded an XH-domain protein whereas *T. oleivorans* contained a Nif11 domain protein (also in Clade 3 genomes) and a downstream region with a VapBC toxin-antitoxin system, a transposase, universal stress proteins, and enzymes associated with detoxification and carbon and peptide metabolism.

### Hydrocarbon-degradation potential

Aliphatic alkane degradation-associated markers were widespread across the 74 genomes regardless of diazotrophic capacity (Supplementary Fig. 9). At least one marker assigned to terminal/biterminal alkane oxidation occurred in all genomes, with the alkane monooxygenase *alkB* in only 7 putative diazotrophs (Clade 3 and Clade 4) compared with 47 non-diazotrophs, and *almA* in one Clade 3 genome. Their electron-transfer genes encoding rubredoxin and rubredoxin reductase occurred broadly in 72 and 67 genomes, respectively. The Finnerty pathway and hydrocarbon uptake-associated genes occurred in 73 and 71 genomes, respectively. Bioplastic degradation markers were also widespread; PBAT-associated esterase occurred in all 25 putative diazotrophs and 46 non-diazotrophs, while PHA/PHB degradation genes occurred in 19 putative diazotrophs comprising all Clade 1, Clade 2, and Clade 4 genomes and in 44 non-diazotrophs. PEG and PCL/PET associated genes occurred in only two non-diazotrophic genomes. Seven genomes contained auxiliary alcohol and aldehyde-dehydrogenase genes including one Clade 4 putative diazotroph (*T. oleivorans*) and 6 non-diazotrophs. In contrast, aromatic hydrocarbon degradation capacity showed pronounced clade-level variation among putative diazotrophs. At least one marker occurred in 17 of 25 genomes comprising all six Clade 1 and all 11 Clade 2 genomes, but none of the Clade 3 or Clade 4 genomes. Clade 1 contained the richest repertoire with multiple pathways (catechol, phenylacetate, toluene, protocatechuate, phenol, benzoate), whereas Clade 2 genomes contained catechol- and phenylacetate-associated markers. Among the 49 non-diazotrophs, six contained aromatic markers. *Thalassolituus* sp. 3um A1 had the broadest repertoire with multiple aromatic degradation pathways, two genomes contained ten markers each, whereas the remaining three contained only one catechol-associated marker.

## Discussion

Biological nitrogen fixation is fundamental to the marine nitrogen cycle, yet how this trait evolved and spread across heterotrophic marine bacteria remains a critical gap in our understanding. Recent characterization of N_2_ fixation in *T. haligoni* raises important questions about the distribution and evolutionary dynamics of *nif* genes within a genus best known for several obligate hydrocarbonoclastic species that dominate oil-contaminated environments (Yakimov et al. 2004; Dong et al. 2022). In the Tara Oceans dataset, exact matches to representative *Thalassolituus* NifH protein sequences were detected across multiple ocean basins and depth layers, although their distributions differed among clades (Fig. 1). Our analysis of key *nif* genes in 74 *Thalassolituus* genomes reveals that N_2_ fixation is restricted to a subset of lineages and shows phylogenetic and compositional signatures that support repeated lateral acquisition of *nif* loci followed by lineage-specific retention and subsequent exchange among closely related diazotrophs in nitrogen-limited and hydrocarbon-rich environments. These findings support a reticulate evolutionary history of N_2_ fixation in *Thalassolituus* and illustrate how diazotrophy can spread within marine heterotrophic microbial communities.

A complete suite of N_2_-fixation genes was observed in only 25 of 74 *Thalassolituus* genomes (33.8%), with the remaining 49 lacking all six core *nifHDKENB* genes (Fig. 2). Most genomes in our dataset (58/74) were MAGs, which can have incomplete gene recovery and may contain contamination (Bowers et al. 2017). The absence of *nif* genes in many *Thalassolituus* genomes is unlikely attributable to assembly artifacts, since MAGs were quality filtered (>85% completeness and <5% contamination) and complete *nif* clusters were consistently recovered in both MAGs and isolate genomes (Supplementary Table 2). Many genomes lacking *nif* were high-quality single-contig isolate assemblies, including several *T. hydrocarbonoclasticus and T. oleivorans* strains, supporting the interpretation that *nif* gene presence and absence reflect true biological variation. Genomes lacking *nifHDKENB* showed only sparse, low-identity hits to regulatory genes *nifA*/*nifL*, and to alternative nitrogenase-associated genes (*anfA*, *vnfA*), which are typically interpreted as divergent, non-diazotrophic homologs rather than evidence of functional N2 fixation (Dos Santos et al. 2012; Koirala and Brözel 2021).

At the broader family scale, genomes assigned to *Thalassolituus* were interleaved with *Oceanobacter, Bacterioplanes, Bacterioplanoides,* and related genera in the Oceanospirillaceae phylogeny rather than forming a clade (Supplementary Fig. 4). Previous phylogenomic analyses similarly resolved Oceanospirillaceae into five well-supported subgroups and proposed dividing the family into more coherent lineages (Liao et al. 2020). More recently, the family was reorganized as Oceanobacteraceae fam. nov. based on phylogenomic relationships and family-level AAI thresholds (Flores-Félix et al. 2025). The maximum-likelihood species tree based on 74 single-copy core genes places putative diazotrophs in four well-supported clades interspersed among non-diazotrophic relatives (Fig. 3), implying that diazotrophy in *Thalassolituus* does not trace back to a single ancestral lineage. Explaining this patchy distribution solely by vertical inheritance would require many independent gene-loss events. In the species tree, *T. oleivorans* SRR3933287 is the only putative diazotroph nested within a well-supported group of otherwise non-diazotrophic *T. oleivorans* strains (bootstrap = 98%), while MAG485 is recovered as the sister lineage to this group (bootstrap = 100%). If their *nif* loci had been inherited vertically from the most recent common ancestor, their absence from the intervening non-diazotrophic lineages would require multiple independent losses. However, SRR3933287 and MAG485 do not group together in the NifHDKENB tree (Fig. 4A) which instead supports relatively recent lateral acquisition and subsequent recombination of their *nif* loci.

Similarly, large genomic and metagenomic datasets have shown that *nif* genes are widely but sporadically distributed, and often phylogenetically incongruent with host genomes due to repeated gene transfer occurring across phyla (Koirala and Brözel 2021; Delmont et al. 2022; Deng et al. 2025). An analysis of the nitrogen fixation in the genus *Paenibacillus* suggested that the ancestral lineage did not fix nitrogen, and N_2_-fixing strains acquired a *nif* cluster via LGT from a *Frankia*-like donor, followed by subsequent loss in some lineages (Xie et al. 2014).

Phylogenetic incongruence and topology tests provide evidence of LGT in the evolutionary history of nitrogen fixation in *Thalassolituus*. Taxa that formed Clades 1 (*P. penaei,* ARC_Gamma03, and *T. haligoni)* and 4 (*T. oleivorans* and MAG485) in the species tree were separated in the Nif trees. Conversely, the *NifHDKENB* tree grouped the HBD_Gamma08 subclade of Clade 1 with the distantly related Clade 3 genomes (Fig. 4A, B). Clade 2 provided a notable contrast because its placement was congruent in the species and Nif trees, which suggests that this lineage potentially inherited the *nif* locus vertically after an earlier acquisition. Studies of other diazotrophs have reported strong concordance among *NifHDKENB* phylogenies, indicating that these proteins co-evolved as a conserved functional module (Koirala and Brözel 2021; Nichio et al. 2025). However, several genomes in our analysis occupied different positions among the individual protein trees, which indicates distinct evolutionary histories and possible post-acquisition reshuffling of *nif* loci (See Supplementary Fig. 5). The Approximately Unbiased (AU) tests provided statistical support for these phylogenetic conflicts; most individual loci and concatenated NifHDK */* NifHDKENB alignments rejected the species tree topology (Table 2).

Although the NifH and NifD alignments did not reject the species tree, their low support likely reflects the limited phylogenetic information in single conserved genes to distinguish among alternative topologies (Poptsova and Gogarten 2007; Palmer et al. 2019). Tree-distance measures also showed moderate rather than complete discordance; this combination supports a mixed evolutionary history in which some lineages likely inherited previously acquired *nif* loci, whereas others acquired them more recently. BLASTP matches (≥95% amino acid identity) and the expanded NifH phylogeny also placed *Thalassolituus* Nif proteins within a broader pool of marine Gammaproteobacteria (Supplementary Table 3; Supplementary Fig. 7). These affiliations varied among clades; ARC_Gamma03 and *T. haligoni* sequences from Clade 1 clustered with *Oceanobacter* and *Halioxenophilus*, Clade 3 grouped with *Marinobacterium ramblicola* and *M. zhoushanense,* and some Clade 2 and Clade 4 sequences clustered together despite their separation in the species tree. Identical NifH sequences also occurred within Clades 1 and 2.

These lineages may represent possible sources or recipients of *Thalassolituus nif* genes, although sequence similarity cannot establish the direction of transfer.

Whereas these broader affiliations identify possible exchange partners, homologous recombination analysis provided evidence of genetic exchange among *Thalassolituus* lineages. Recombination can also generate discordant gene histories and likely contributes to the low phylogenetic resolution we see in the NifH and NifD protein trees (Shikov et al. 2022). Eleven well-supported recombination events were detected across *nif* loci in *Thalassolituus,* in which the same genomes appeared as both recombinants and parents, including *T. oleivorans* SRR3933287 and MAG485 (Clade 4), and *T. haligoni and P. penaei* (Clade 1) (Supplementary Table 6; Fig. 5). These non-random signals in genomes that also show the strongest conflicts with the species tree are unlikely to be coincidental and help explain why their placement varies even among individual Nif protein trees. In the *nifH* gene, multiple recombination events involving *T. oleivorans* and MAG485 span almost the entire nucleotide sequence and suggest ongoing genetic exchange among these lineages. Similar recombination-driven rearrangements of *nif* genes have been documented in other diazotrophs, including fragmentation of *nifBHDK* into thirteen fragments spread over 350 kb in the cyanobacterial diazotroph *Calothrix* sp. NIES-4101 (Hirose et al. 2021), and large deletions or rearrangements mediated by insertion sequences in *Bradyrhizobium* (Arashida et al. 2022).

If different *Thalassolituus* lineages acquired *nif* genes through LGT, their genomic neighborhoods should retain signatures of integration and subsequent divergence, as seen for laterally acquired *nif* clusters in other bacteria (Nichio et al. 2025). Across *Thalassolituus*, the *nifHDK* genes form a conserved syntenic block (Fig. 6), while *nifE* and *nifN* remained directly adjacent despite variation in their distance from *nifHDK.* This conservation likely reflects selection to maintain coordinated expression and interactions among the subunits (Raymond et al. 2004). In contrast, the surrounding regions varied among clades. Molybdate transporter genes (*modABC*) occurred downstream of *nifHDK* in 17 genomes and may represent supportive elements that have coevolved or were corecruited with *nif* to supply molybdenum for FeMo-cofactor biosynthesis (Delgado et al. 2006; Pi et al. 2025). Flanking monothiol glutaredoxins and ferredoxin-associated proteins may also support nitrogenase maturation through biogenesis of iron-sulfur clusters and oxidative stress protection. SIR2-2 domain proteins may regulate N_2_ fixation post-translationally through lysine deacetylation, as observed in *Zymomonas mobilis* (Nisar et al. 2021). Ion-translocating oxidoreductase complexes found in some neighborhoods could contribute to the supply of low-potential electrons required by nitrogenase (Zhang and Einsle 2024). Several consistently flanking domain-containing proteins may also support N_2_ fixation although most of their biochemical functions remain unknown. Nif11 associated with *nif* loci, for example, was originally described in the diazotroph *Azotobacter vinelandii* yet its function remains ambiguous, and related domains also occur in precursor peptides for post-translationally modified products (Haft et al. 2010). Ankyrin-repeat proteins in 10 genomes, which mediate protein-protein interactions and are most common in eukaryotes, have been proposed to act as mobile modules transferred laterally across distantly related taxa (Bork 1993).

The conserved accessory structure of Clade 2 supports stable inheritance of an integrated *nif* region, and the greater heterogeneity of Clades 1 and 4 may reflect signatures of recombination after acquisition. Similar variation in *nif* gene organization and highly variable accessory and regulatory gene content occurs across other diazotrophs and may reflect lineage-specific LGT and ecological adaptation (Setubal et al. 2009; Xie et al. 2014; Tao et al. 2021).

Several recombination and repair-associated genes also occurred near *nifHDK*, including DNA topoisomerase III in Clade 1, exodeoxyribonuclease VII (*xseB*) in Clade 2, and DEAD/DEAH-box and RecG helicases in HBD_Gamma08. Mobile genetic elements (MGEs) are commonly identified by hallmark mobility genes that can leave signatures of recent acquisition, including integrases or recombinases, transposases, phage-associated genes, and nearby tRNA integration sites (Khedkar et al. 2022). These signatures were unevenly distributed although HBD_Gamma08 (Clade 1) contained the strongest evidence of a laterally acquired region. Its *nifHDK* region contained a nearby phage-tail protein and a larger putative mobile region upstream with an integrase, tRNA genes, and several phage-associated functions. These features are characteristic of a putative genomic island (Juhas et al. 2009). Numerous hypothetical proteins flanking the integrase to the end of the contig suggest that the element may extend beyond the annotated mobile region. A Rop-family plasmid RNA-binding protein was also located 1 kb downstream of *nifEN*, which provides a possible signature of plasmid-associated mobility. In *T. oleivorans*, a transposase and VapB/VapC toxin-antitoxin pair occurred next to the *nif* neighborhood. Type II toxin-antitoxin systems often function as addiction modules that stabilize mobile elements and large genomic regions (Fraikin et al. 2020); their co-occurrence with a transposase near *nifHDK* supports the possibility that this region was laterally acquired. HBD_Gamma04 and HBD_Gamma06 similarly contained isolated phage-tail proteins near *nifHDK* but lacked additional MGEs. The absence of comparable mobility genes from other neighborhoods does not exclude older acquisitions because genomic islands can lose these genes and gradually acquire the sequence composition of their host, and fragmented MAGs may not recover complete genomic island boundaries.

Codon usage bias is the non-random use of synonymous codons and reflects the combined action of mutation, selection, and drift on coding sequences (Iriarte et al. 2021). Because laterally acquired genes often arrive with donor-like codon usage that deviates from host preferences, atypical codon usage provides a useful complementary signal for identifying candidate transfers (Sharp and Li 1987; Iriarte et al. 2021). Across *Thalassolituus*, *nifHDK* genes show higher preference for host-preferred codons than the biosynthesis *nifENB* genes, with *nifH* generally the most optimized subunit (Supplementary Table 5; Supplementary Fig. 8). This is likely due to highly expressed, functionally critical genes that tend to evolve stronger codon bias toward abundant tRNAs as this adaptation preserves proteome-wide translation efficiency (Frumkin et al. 2018). The most informative signal emerged from *nifH* codon adaptation among clades. Clade 2 genomes, which were congruent between the species and Nif phylogenies, show the highest *nifH* optimization, suggesting long-term integration of *nif* systems and gradual amelioration toward host codon preferences (Callens et al. 2021). Genomes with the strongest phylogenetic conflicts and recombination signals show substantially lower *nif* CAI values, including the Clade 4 singletons and *P. penaei* from Clade 1, indicating weaker adaptation to host-preferred codons, which is compatible with more recent acquisition but may also reflect differences in expression and selective constraints. Similar patterns were reported by Mondal et al. (2016), who found that GC-driven mutational bias and translational selection together shape codon usage in *nifHDK* genes across phylogenetically diverse diazotrophs.

Beyond the evolutionary history of *nif*, *Thalassolituus* fits the classical profile of obligate hydrocarbonoclastic bacteria and has a near-universal capacity for aliphatic alkane degradation (Supplementary Fig. 9). Genomic and proteomic studies show that members of the genus rely largely on petroleum-derived compounds as carbon and energy sources (Yakimov et al. 2004; Gregson et al. 2018; Dong et al. 2022). Aromatic hydrocarbon capacity, however, varied among the putative diazotrophs. Clade 1 contained the broadest range of pathways, including catechol, phenylacetate, toluene, protocatechuate, phenol, and benzoate utilization, whereas Clade 2 mostly retained the phenylacetate pathway, and Clades 3 and 4 showed no aromatic hydrocarbon capacity. Clade 1 may therefore have the greatest potential to combine diazotrophy with aromatic hydrocarbon utilization. Diazotrophy may provide an ecological advantage and could sustain growth when fixed nitrogen becomes depleted during hydrocarbon degradation (Shin et al. 2019), although aerobic hydrocarbon metabolism and oxygen-sensitive nitrogenase require different conditions. Surface oil slicks at the air-water interface illustrate this constraint because oxygen exposure supports hydrocarbon oxidation but may restrict N_2_ fixation (Voskuhl and Rahlff 2022). *T. haligoni* BB40 is the only cultured diazotrophic *Thalassolituus* studied physiologically. It showed low fixation activity when fixed nitrogen was available, but growth with N_2_ as the sole nitrogen source occurred only under hypoxia where fixation rates were much higher (Rose et al. 2024; Rose et al. 2026). Evidence from other systems supports this; hydrocarbonoclastic species carry *nif* genes and exhibit nitrogenase activity (Musat et al. 2006; Dashti et al. 2015), and metagenomic analyses show significant co-occurrence between *nifH* and PAH-degradation genes (Zhang et al. 2019). *Polaromonas naphthalenivorans* reportedly acquired *nif* via LGT and increases its expression in response to hydrocarbons (Foght 2010), whereas *Candidatus* Macondimonas diazotrophica encodes both functions and is abundant in oil- impacted coastal sediments (Karthikeyan et al. 2019).

Our results show that diazotrophy is restricted to a subset of *Thalassolituus* lineages, and that together with other key findings, the distribution and diversification of *nif* genes is best explained jointly by lateral gene transfer, vertical inheritance, homologous recombination, and lineage-specific retention. Some *nif* clusters, particularly those in Clade 2, appear to have become stably integrated and subsequently inherited with their host lineages, whereas the greater phylogenetic discordance, recombination, and genomic signatures observed in Clades 1 and 4 suggest more recent or ongoing genetic exchange. Although closely related Gammaproteobacteria represent possible participants in this exchange, the direction and source of transfer cannot be determined from sequence similarity alone. These evolutionary patterns appear closely linked to ecology, as facultative anaerobic and diazotrophic *Thalassolituus* associated with hydrocarbon-rich environments are impacted by nitrogen limitation. The co- occurrence of nitrogen fixation and hydrocarbon utilization genes further identifies a potential connection between diazotrophy and hydrocarbon-associated lifestyles; diazotrophy may support growth when fixed nitrogen becomes limiting in hydrocarbon-rich environments. Future studies combining cultivation, expression analyses, proteomics, and direct measurements of N_2_ fixation are needed to determine when these pathways are active and to assess the contribution of heterotrophic bacterial diazotrophs to nitrogen cycling in hydrocarbon-rich marine ecosystems. More broadly, our findings demonstrate how the evolutionary histories of complex metabolic traits can differ from those of their microbial hosts and establish *Thalassolituus* as an important lineage for investigating the acquisition and integration of N_2_ fixation in marine bacteria.

## Acknowledgements

This work was supported by the Faculty of Computer Science and the Department of Biology at Dalhousie University, and funding was provided by the Natural Sciences and Engineering Research Council of Canada (NSERC) to Robert G. Beiko and Julie LaRoche. Computational resources were provided by the Digital Research Alliance of Canada. The authors thank John Archibald and Erin Bertrand for their helpful comments and insights on earlier versions of this work. ChatGPT version 5 was used to assist in drafting and debugging R code used to generate selected visualizations. All resulting helper scripts and outputs were modified and verified by the authors.

## Author contributions

**SSB:** conceptualization, data curation, formal analysis, investigation, methodology, software, validation, visualization, writing – original draft, writing – review and editing. **RGB and JL:** conceptualization, funding acquisition, methodology, project administration, resources, supervision, validation, writing – review and editing. All authors read and approved the final manuscript.

## Data availability

All genomic data and associated metadata used in this study are publicly available through the NCBI database, with all accession numbers provided in Supplementary Tables 1 and 2. Custom scripts and selected data files are currently stored in a private GitHub repository and are available upon request. The repository will be made public upon publication of the peer- reviewed article.

