## Supplementary Material for "Lateral gene transfer shapes the distribution of nitrogen fixation within a cosmopolitan clade of marine *Thalassolituus*"

Supplementary Tables S1-S6 are available in the project GitHub repository: <https://github.com/Soma-barawi/Proj-Thalassolituus>.

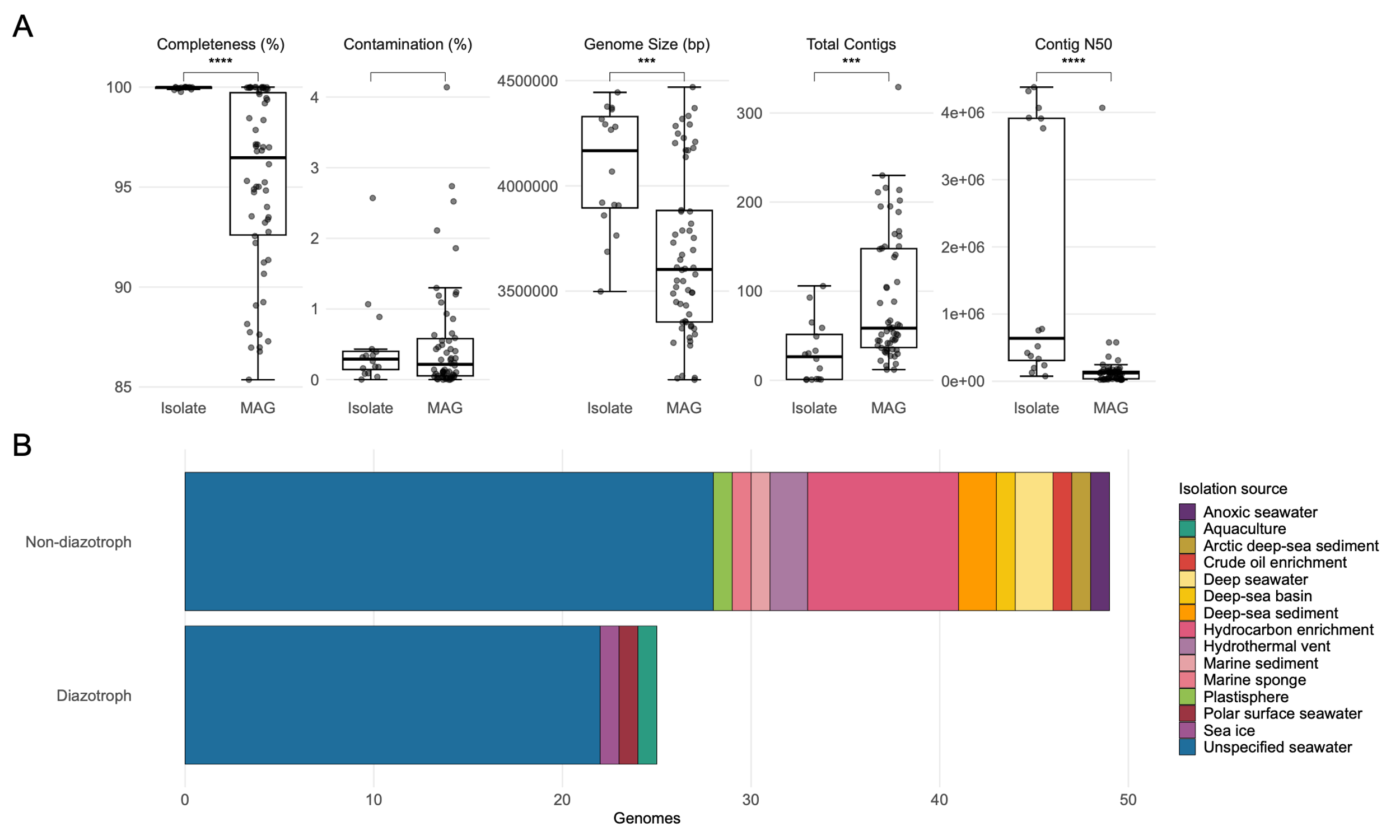

**Supplementary Figure 1.** Genome quality metrics and isolation sources of 74 *Thalassolituus* genomes. (A) Boxplots show CheckM2 completeness and contamination, genome size, number of contigs, and contig N50 values across isolate genomes and MAGs (metagenome-assembled genomes). The central lines indicate the median, boxes span the interquartile range (IQR), and whiskers extend to data points within 1.5 times the IQR. Statistical significance was assessed using two-sided Wilcoxon rank-sum tests with Benjamini-Hochberg FDR-adjusted P values (**P ≤ 0.01, and ***P ≤ 0.001). (B) Isolation sources among putative diazotrophic and non-diazotrophic genomes.

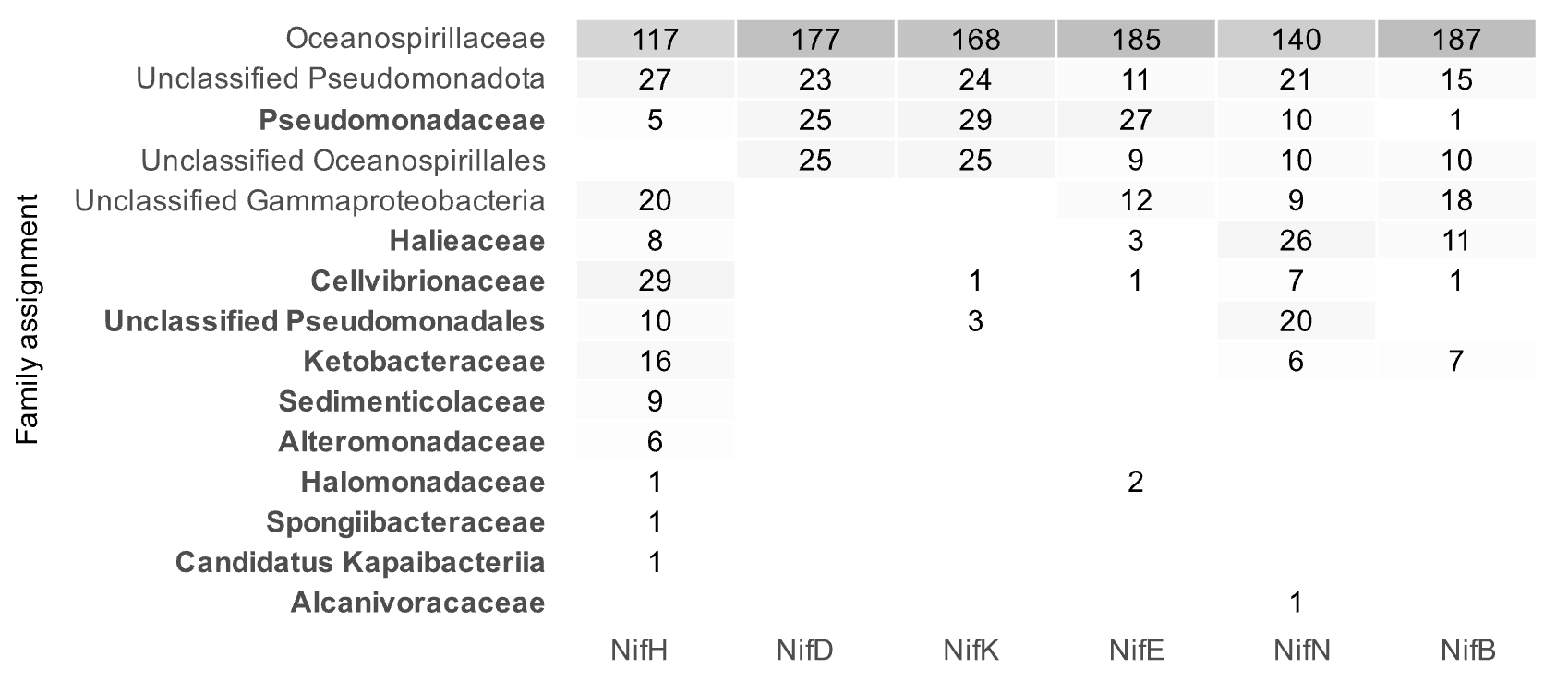

**Supplementary Figure 2**. Summary of BLASTP analysis of Nif proteins in *Thalassolituus*. Family level classification of the top 10 homologous hits for each Nif protein across the 25 putative diazotrophic genomes. Unclassified labels indicate the closest taxonomic rank available when family level classification was not assigned. Assignments outside the expected *Thalassolituus* lineage are highlighted in bold.

**Supplementary Figure 3**. Pairwise comparisons of (A) Average Nucleotide Identity (ANI) and (B) Average Amino Acid Identity (AAI) between 74 *Thalassolituus* genomes. The 95% ANI and 65% AAI thresholds represent species and genus-level delineations. The 25 putative diazotrophs are labeled in red.

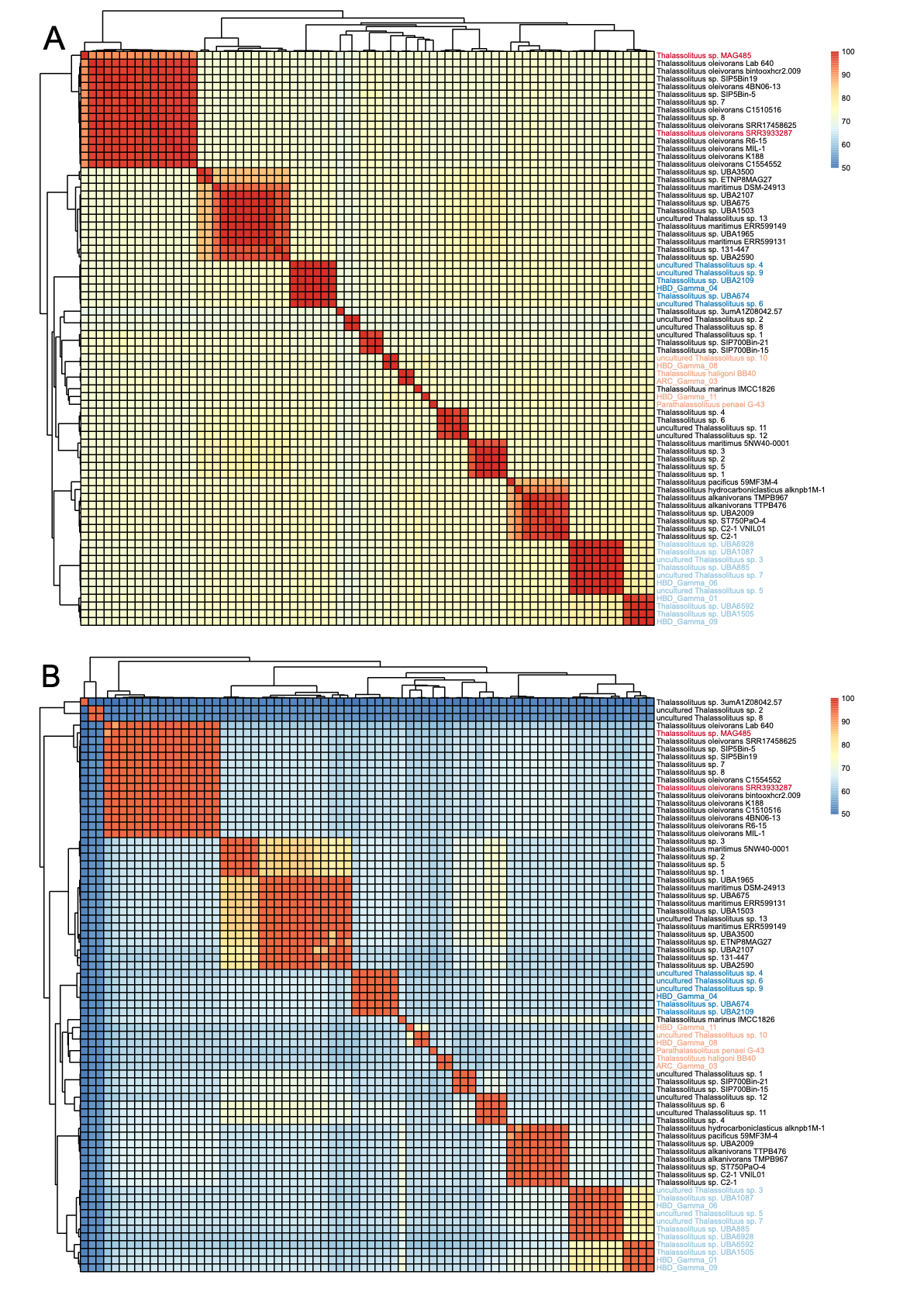

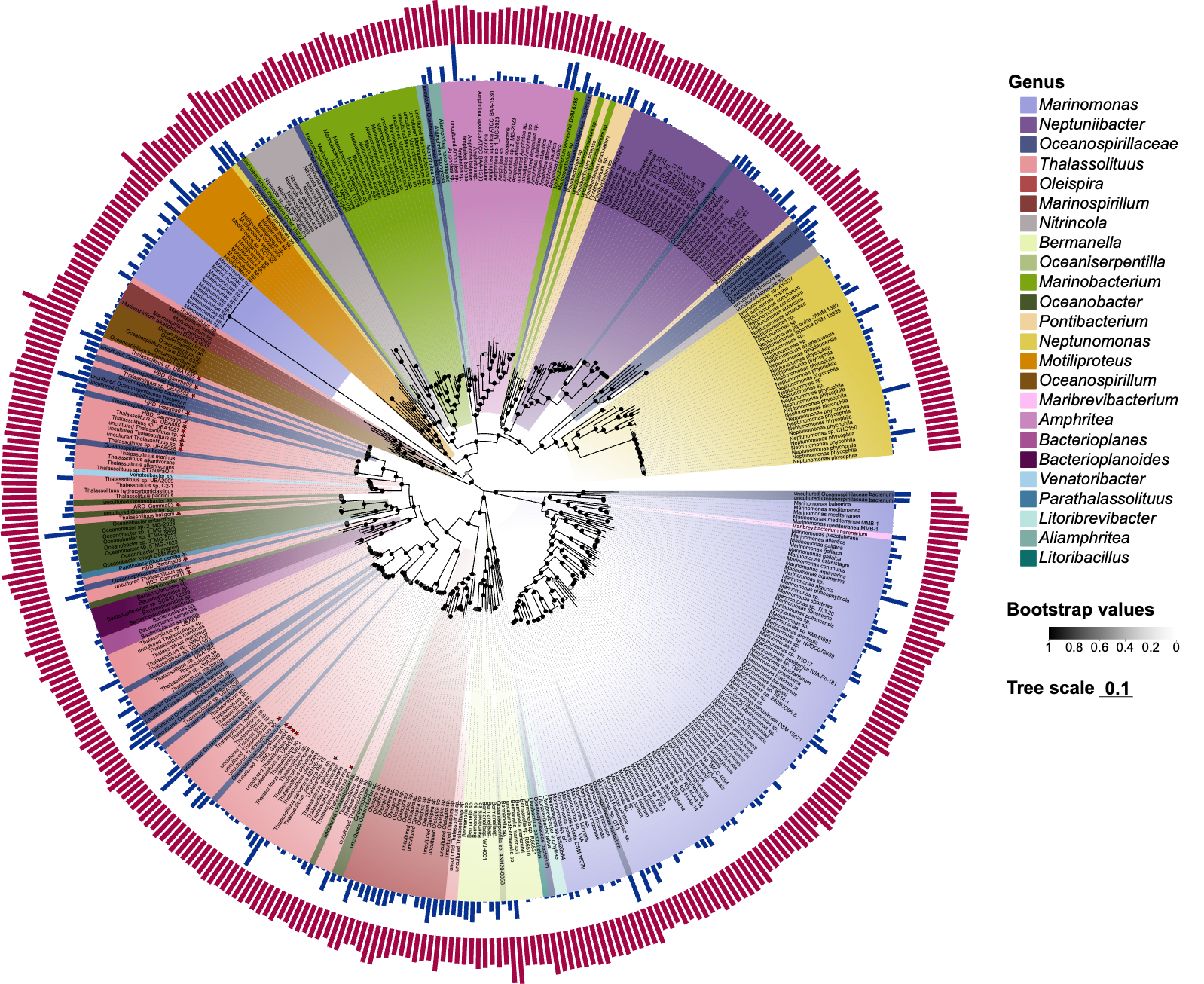

**Supplementary Figure 4.** Phylogeny of Oceanospirillaceae. Maximum-likelihood unrooted tree of 421 high-quality genomes reconstructed from 120 “soft-core” genes identified via Panaroo and present in 95% to 99% of the genomes across the dataset. Bootstrap support values are indicated at nodes with black circles >90% and gray circles 70-90%. The scale bar represents the mean number of substitutions per site.

**Supplementary Figure 5**. Phylogenies of Nif proteins. Maximum-likelihood trees for NifH, NifD, NifK, NifE, NifN, and NifB proteins identified in 25 putative diazotrophic *Thalassolituus* genomes.  Bootstrap support values are shown at the nodes. Scale bars represent the mean number of substitutions per site. All trees are rooted using the corresponding homologous sequences from *Bradyrhizobium diazoefficiens* USDA 110.

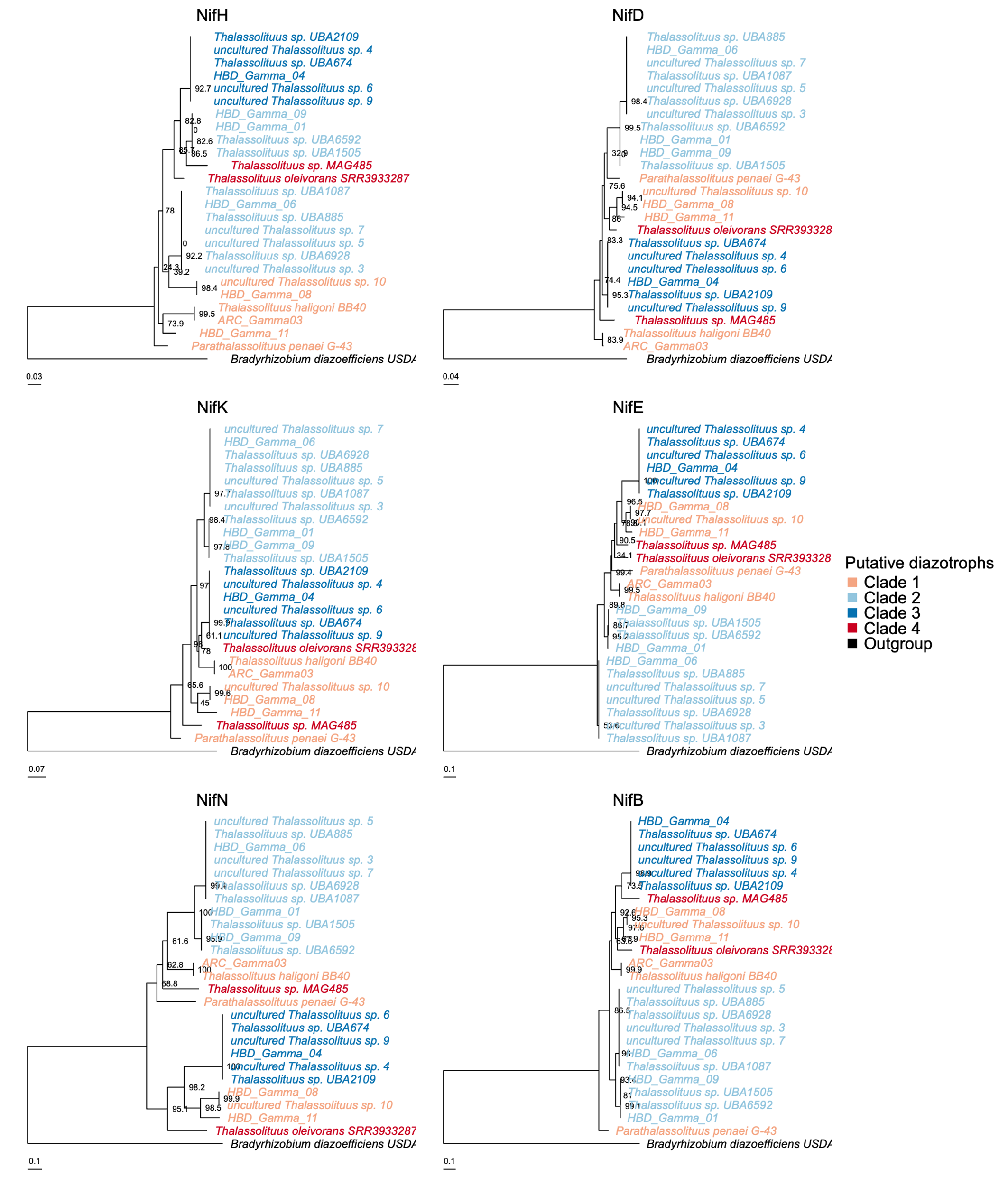

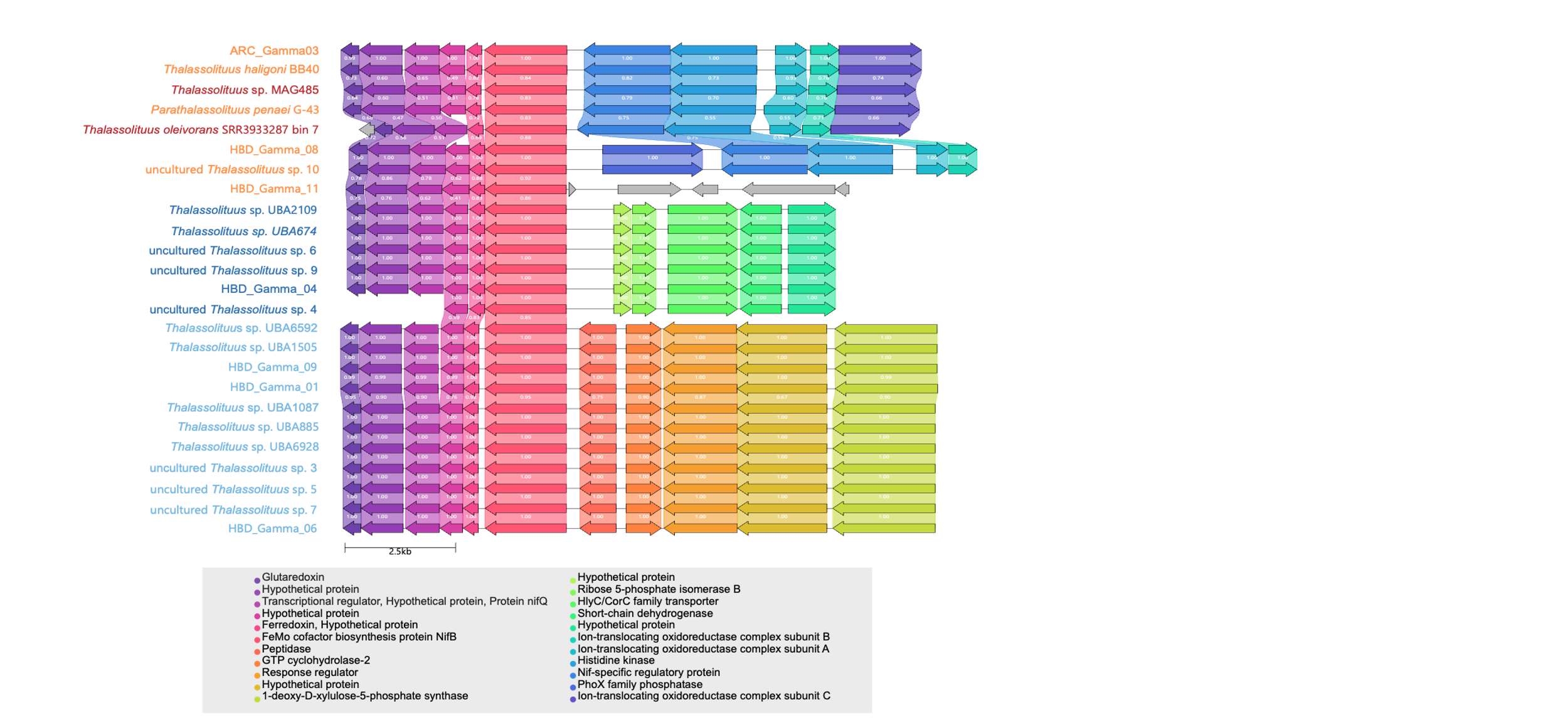

**Supplementary Figure 6.** Genomic neighborhoods of *nifB* and ten flanking genes in 25 putative diazotrophic *Thalassolituus* genomes. Genes are drawn to scale as arrows with their orientation indicating the direction of transcription. Colors denote homologous gene groups identified by global pairwise alignments, and connecting links represent the degree of amino acid sequence identity.

**Supplementary Figure 7.** Expanded phylogeny of NifH sequences. (A) Maximum-likelihood tree of 4550 NifH sequences from the NCBI RefSeq database and (B) subtree highlighting the placement of *Thalassolituus* NifH sequences relative to neighboring diazotrophs. Bootstrap support values are indicated at nodes with black circles >90% and gray circles 70-90%. The scale bar represents the mean number of substitutions per site.

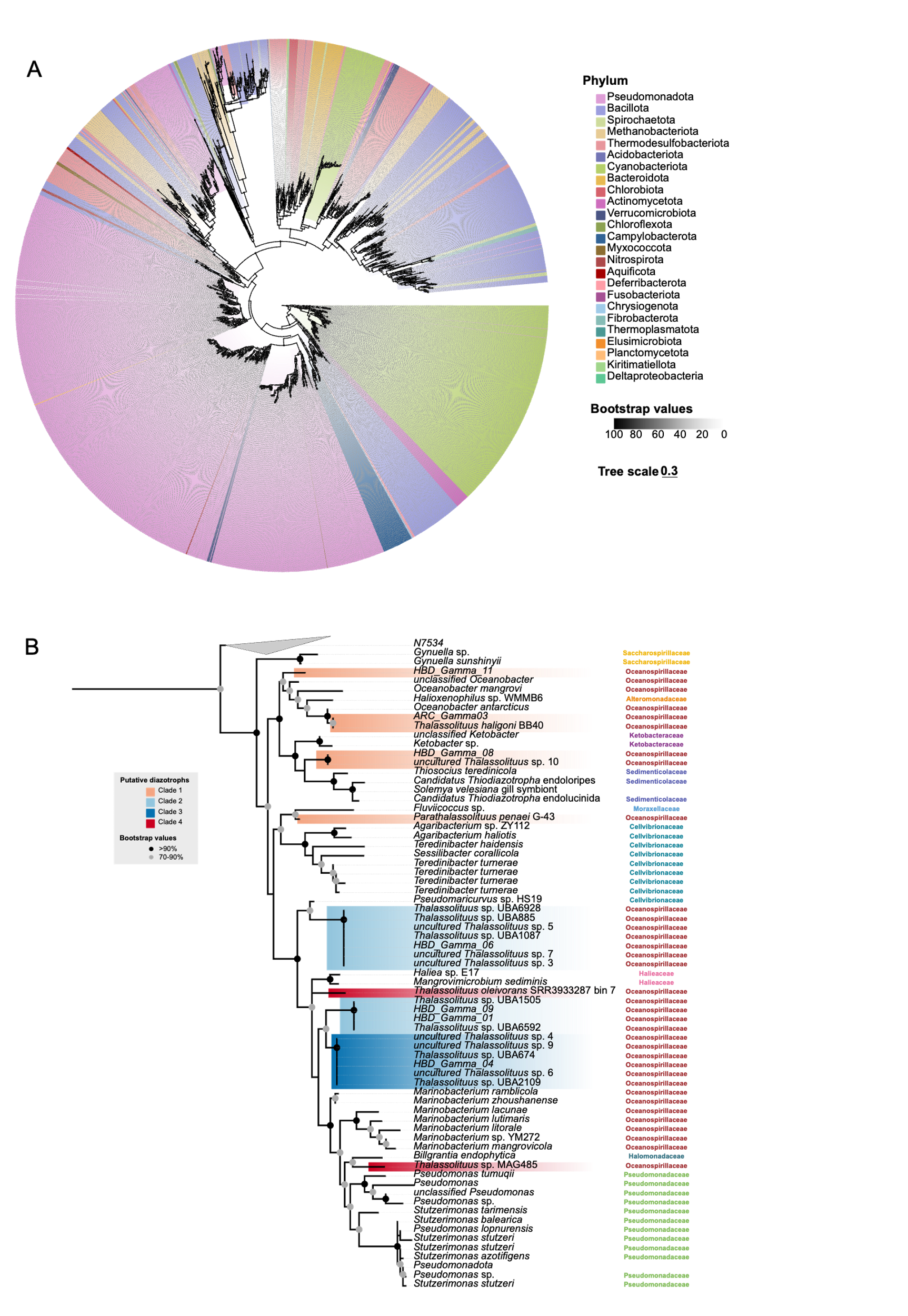

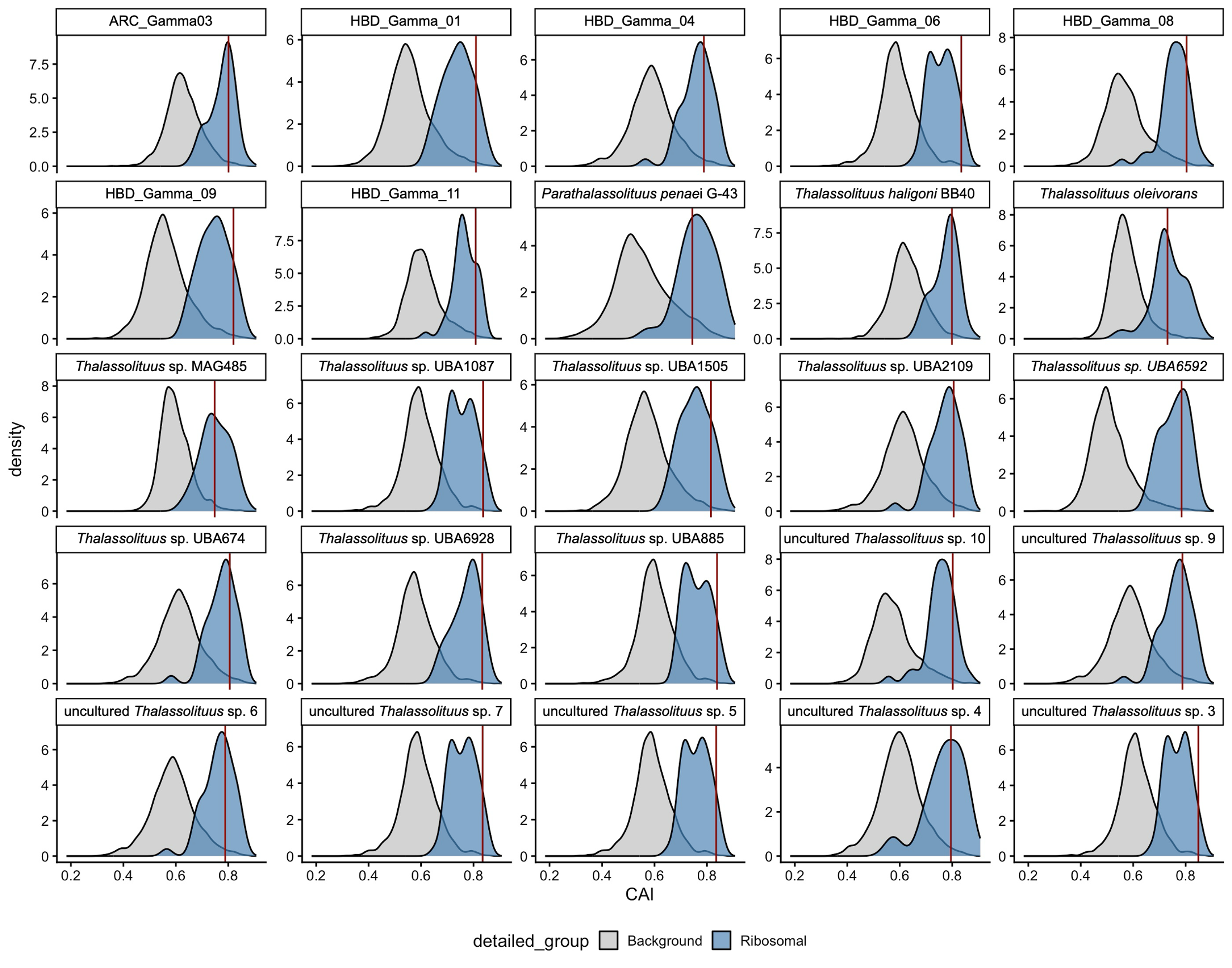

**Supplementary Figure 8.** Codon adaptation index (CAI) distributions across 25 putative diazotrophic *Thalassolituus* genomes. Each density plot shows the distribution of CAI values for all background coding DNA sequences alongside the genome-specific reference set of highly expressed ribosomal genes. The vertical red line indicates the specific CAI value of *nifH* within each genome.

**Supplementary Figure 9**. Distribution of hydrocarbon degradation pathways across 74 *Thalassolituus* genomes. The bubble plot shows the abundance of predicted genes identified using the Hydrocarbon Aerobic Degradation Enzymes and Genes (HADEG) database, with bubble size corresponding to the number of gene hits predicted for each specific subpathway.

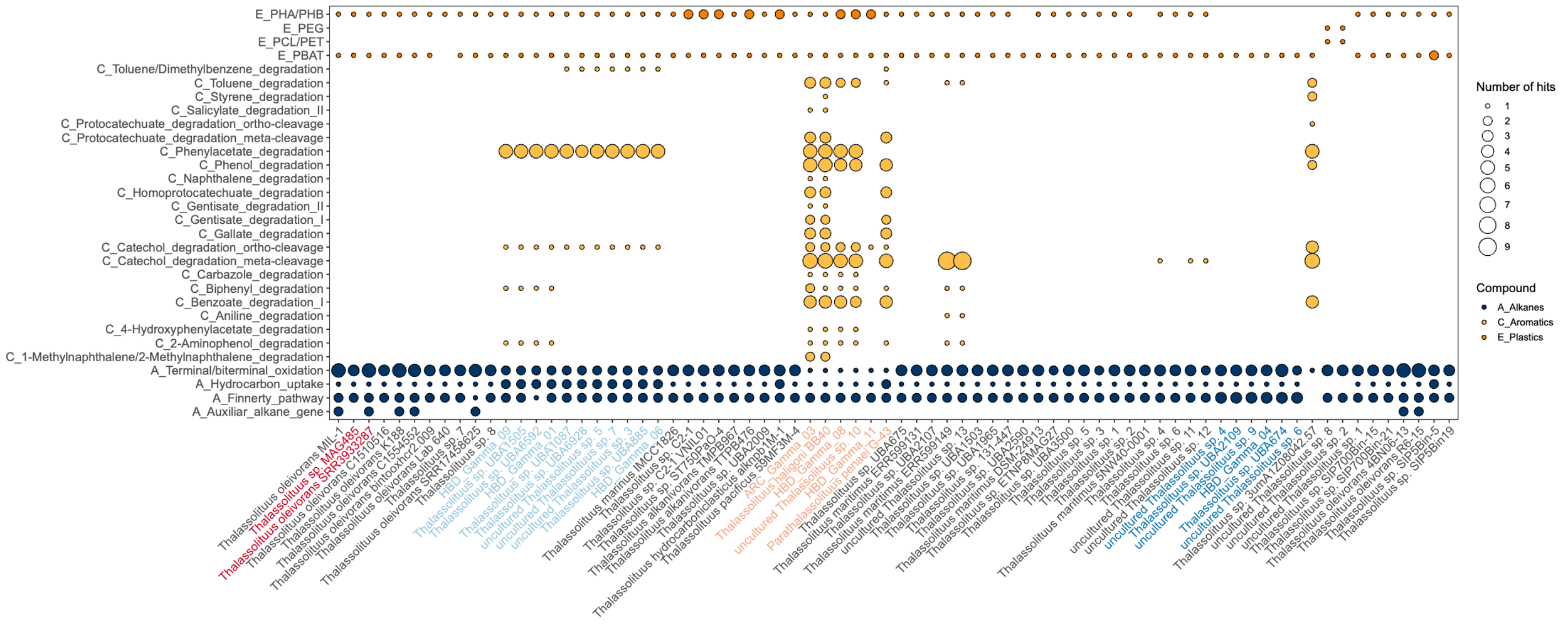
